# Dynamically Evolving Cell Sizes During Early Development Enable Normal Gastrulation Movements In Zebrafish Embryos

**DOI:** 10.1101/481325

**Authors:** Triveni Menon, Asfa Sabrin Borbora, Rahul Kumar, Sreelaja Nair

**Affiliations:** Department of Biological Sciences, Tata Institute of Fundamental Research, Homi Bhabha Road, Colaba, Mumbai, 400005, India

**Author notes:** Corresponding author: Sreelaja Nair.

**Keywords:** Zebrafish, cell size, gastrulation, cell migration

## Abstract

Current knowledge of the mechanisms of cell migration is based on differentiated cells in culture where it is known that the actomyosin machinery drives migration via dynamic interactions with the extracellular matrix and adhesion complexes. However, unlike differentiated cells, cells in early metazoan embryos must also dynamically change cell sizes as they migrate. The relevance of cell size to cell migration and embryonic development is not known. Here we investigate this phenomena in zebrafish embryos, a model system in which reductive cell divisions causes cell sizes to decrease naturally over time as cells migrate collectively to sculpt the embryonic body plan. We show that cell size reduction during early development follows power-law scaling. Because mutations that can perturb cell sizes so early in development do not exist, we generate haploid and tetraploid zebrafish embryos and show that cell sizes in such embryos are smaller and larger than the diploid norm, respectively. Cells in embryos made of smaller or larger than normal cells migrate sub-optimally, leading to gastrulation defects. Multiple lines of evidence suggest that the observed defects originate from altered cell size rather than from pleotropic effects of altered ploidy. This interpretation is strengthened by the result wherein restoring cell sizes to normal diploid-like values rescues gastrulation defects. Live imaging of chimeric embryos where haploid/tetraploid cells are introduced into diploid embryos reveal the cell-autonomous nature of the migration defects. Additionally, aberrant intracellular actin dynamics with respect to the vectorial direction of motion suggests a cellular mechanism behind the migration defects. Taken together, early reductive cell divisions potentially allow dynamic, stage-specific cell size norms to emerge, which enables efficient collective cell migration to correctly position cells in space and time to shape an amorphous ball of blastoderm into an embryo.

## INTRODUCTION

Cell migration is a complex interplay between actomyosin contractility in migrating cells, the molecular nature of the communication between cells and their environments, the physical properties of the extracellular environment and physical contacts between neighboring cells (Lee & Losert, 2019, Martin & Lewis, 1992, Norden & Lecaudey, 2019, Petrie, Gavara et al., 2012, Schaks, Giannone et al., 2019, Trepat, Wasserman et al., 2009). Cell migration is also an integral feature of both natural and disease conditions, such as embryonic development, cancer metastasis and wound healing. Though the molecular and cellular mechanisms that drive cell migration are known, how sizes of cells within a migrating collective would influence migration is not known. This is because the molecular and cellular basis of cell migration has been largely gleaned from cells in culture or from differentiated cells later in embryonic development (Lee & Losert, 2019, Norden & Lecaudey, 2019, Petrie et al., 2012, Schaks et al., 2019, Trepat et al., 2009).

Cell size is a fundamental feature of biological systems, which varies across organisms and across cell types within a multicellular organism (Arendt, 2007, Levy & Heald, 2015). In unicellular eukaryotes such as yeast, optimal cell size is an integrated property that emerges as a balance between cell growth and cell division (Umen, 2005). In metazoans, cell size must be viewed minimally in two diverse contexts: adult homeostasis and embryonic development. During homeostasis, organs in the adult animal and the organism as a whole, strives to maintain an optimal organ/tissue size by balancing cell proliferation with cell growth. This balancing act during homeostasis is heavily influenced by autonomous and non-autonomous cues of growth factor signaling (Artiles, Anastasia et al., 2009, Conlon & Raff, 1999, Edgar, 2006, Jorgensen, Nishikawa et al., 2002, Jorgensen & Tyers, 2004, Nurse, 1985, Tumaneng, Russell et al., 2012). The phenomena of cell growth and division can be coupled to or uncoupled from each other in diverse developmental and homeostasis contexts (Dong, Feldmann et al., 2007, Qu, Weiss et al., 2004, Su & O’Farrell, 1998). In general, blocking cell division leads to cell growth, whereas blocking cell growth elicits a cell division arrest (Jorgensen & Tyers, 2004). An exception to this general rule is the early developmental phase of all metazoan embryos wherein the absence of cell growth does not evoke a cell division arrest. On the contrary, early embryonic development in all metazoans is characterized by a visually striking phenomenon of progressive reduction in cell size precisely because cell division is uncoupled from cell growth. This allows newly fertilized single-cell zygotes to generate increasing numbers of progressively smaller daughter cells to form a multicellular embryo. Thus, the inherent nature of early cell divisions in metazoan embryos generates an ever-changing cell size landscape, on a temporal scale of minutes to hours depending on the species.

Experimental analysis in metazoan embryos reveals that when cell size is constantly changing during reductive cell divisions, sizes of intracellular macromolecular assemblies such as the nucleus and the mitotic spindle scales with cell size (Good, Vahey et al., 2013, Levy & Heald, 2010, Loughlin, Wilbur et al., 2011). Thus, embryos must have mechanisms to sense cell sizes early in development, even when cell size is a constantly evolving parameter with each subsequent division. Cell size control has been studied with respect to whole body size of an organism and, is a complex interplay of cell autonomous and non-autonomous events, which are organism, organ or context specific (Henery, Bard et al., 1992, Lloyd, 2013). In general, individual cell sizes in adult animals remain comparable despite dramatic variations in body size, suggesting that cell numbers, rather than cell size is modulated to achieve a body size (Conlon & Raff, 1999, Day & Lawrence, 2000). This idea has been experimentally tested in developing mammalian and amphibian embryos. Increasing or decreasing blastomere numbers in mouse embryos or fusing two 8-cell mouse embryos into one results in embryos of the normal size, an outcome that is achieved by regulation of cell proliferation during development (Lewis & Rossant, 1982, Rossant, 1976, Tarkowski, 1959). Similarly, tetraploid amphibian embryos are made of larger and fewer cells at early blastula stages, but adjust cell numbers to attain normal larval sizes eventually (Conlon & Raff, 1999, Day & Lawrence, 2000, Gibeaux, Miller et al., 2018). In these experiments, after the initial manipulations, embryo size was the end-point readout and the effect on cell size during early development remained unaddressed. Thus, the fundamental question of the importance of cell sizes in early embryonic development for subsequent normal embryogenesis has remained open. Interestingly, despite extensive mutagenesis screens there have been no mutants in animal models in which early cell sizes during reductive cell divisions are shown to alter. Therefore, it also remains unclear whether cell size norms exist at all during early reductive divisions.

We investigated the relevance of cell sizes during the reductive cell division phase on vertebrate embryonic patterning using the rapidly dividing zebrafish embryo as the experimental paradigm (Kimmel, Ballard et al., 1995). In addition to nutrient and growth factor signaling, genome size is known to determine cell size (Cavalier-Smith, 2005, Lee, Davidson et al., 2009, Orr-Weaver, 2015, Otto, 2007). Therefore, in the absence of mutants, to obtain embryos with cell sizes smaller or larger than the normal diploid embryos, we generated haploid and tetraploid zebrafish embryos. We additionally also altered cell sizes in diploid embryos by transient pharmacological treatments. These independent approaches allowed the experimental generation of zebrafish blastoderms made of cells that were smaller or larger cells than the developmental stage-specific cell size norm. Such embryos developed epiboly and gastrulation defects and were morphologically abnormal. Comparative transcriptome analysis showed that the global transcriptomes as well as levels of candidate transcripts of the planar cell polarity (PCP) pathway remained unperturbed. Further, cell size deviations towards smaller and larger than the norm *both* elicited similar defects in migration and gastrulation indicating that mechanisms beyond transcriptional alterations potentially drive emergence of the phenotypes. Importantly, epiboly and gastrulation defects in haploid embryos, which are composed of cells smaller than the norm, could be rescued by experimentally increasing cell sizes before initiation of gastrulation. We find that smaller or larger than normal cells migrate sub-optimally during gastrulation, follow abnormally tortuous migration paths, and exhibit gross defects in intracellular actin localization as they migrate. Thus, the inability of sub-optimally sized cells to migrate efficiently may explain the epiboly and gastrulation defects that emerge when cell sizes in the zebrafish blastoderm deviates from the norm. Our study suggests that a developmental stage-specific cell size norm exists, and cells sizes are dynamically set with each round of cell division to produce a cell size landscape that ensures optimal cell migration leading to normal embryonic development.

## RESULTS

### Reductive cell divisions lead to power law reduction of cell sizes in the zebrafish blastoderm

Observations of embryonic development show that early cell divisions in newly fertilized zygotes result in daughter cells that are progressive smaller in size. However, no studies have quantified the actual sizes of cells during this phase in any metazoan embryo. We therefore began our investigations by characterizing cell sizes in normal diploid zebrafish embryos (Fig. 1, Supplementary Fig. 1A, D, Supplementary Fig. 2A). We chose the early blastoderm at 64-cell stage (2 hours post fertilization hpf), late blastoderm at 1k-cell (3 hpf) and sphere stages (4 hpf), early gastrula at shield stage (6 hpf) and early larva (24 hpf). Cell sizes decreased progressively during successive stages of embryonic development (Fig. 1, Supplementary Fig. 1A, D and Supplementary Fig. 2A). We quantified cell areas from the deep layer of the blastoderm using the BioVoxxel toolbox in ImageJ (Brocher, 2014) (Fig. 1G-J). The average cell size in a zebrafish embryo decreased ∼200 fold during the first 24 hours of development (Fig. 1K, O). Average cell areas decreased by ∼28 fold during the first 6 hours of development, with a decrease of ∼7 fold occurring between 1k-cell and shield stages (Fig. 1K-O). Regression analysis on a log-log plot revealed a linear reduction of the cell areas between 64-cell to shield stages, thus suggesting a power law reduction in cell size (Size ∼Time^N^), where N = -3 is the calculated slope (Fig. 1P, Q, Supplementary Fig. 4A). To obtain a better understanding of how cell sizes evolve dynamically during development, we binned the cells as small, medium and large based on cell areas as follows: at any stage of development, medium size cells were defined as cells with areas +/-10% of the average area for that stage of development. Cells with areas less than this bin were categorized small and those with areas more than this bin were categorized as large (Supplementary Fig. 3A-D). This size categorization allowed us to estimate potential changes in proportions of cells within a specific size category across developmental stages. Interestingly, despite the heterogeneity in cell sizes, the relative proportions of cells within a particular size category remained comparable across developmental stages (Fig. 1R).

**Figure 1:**
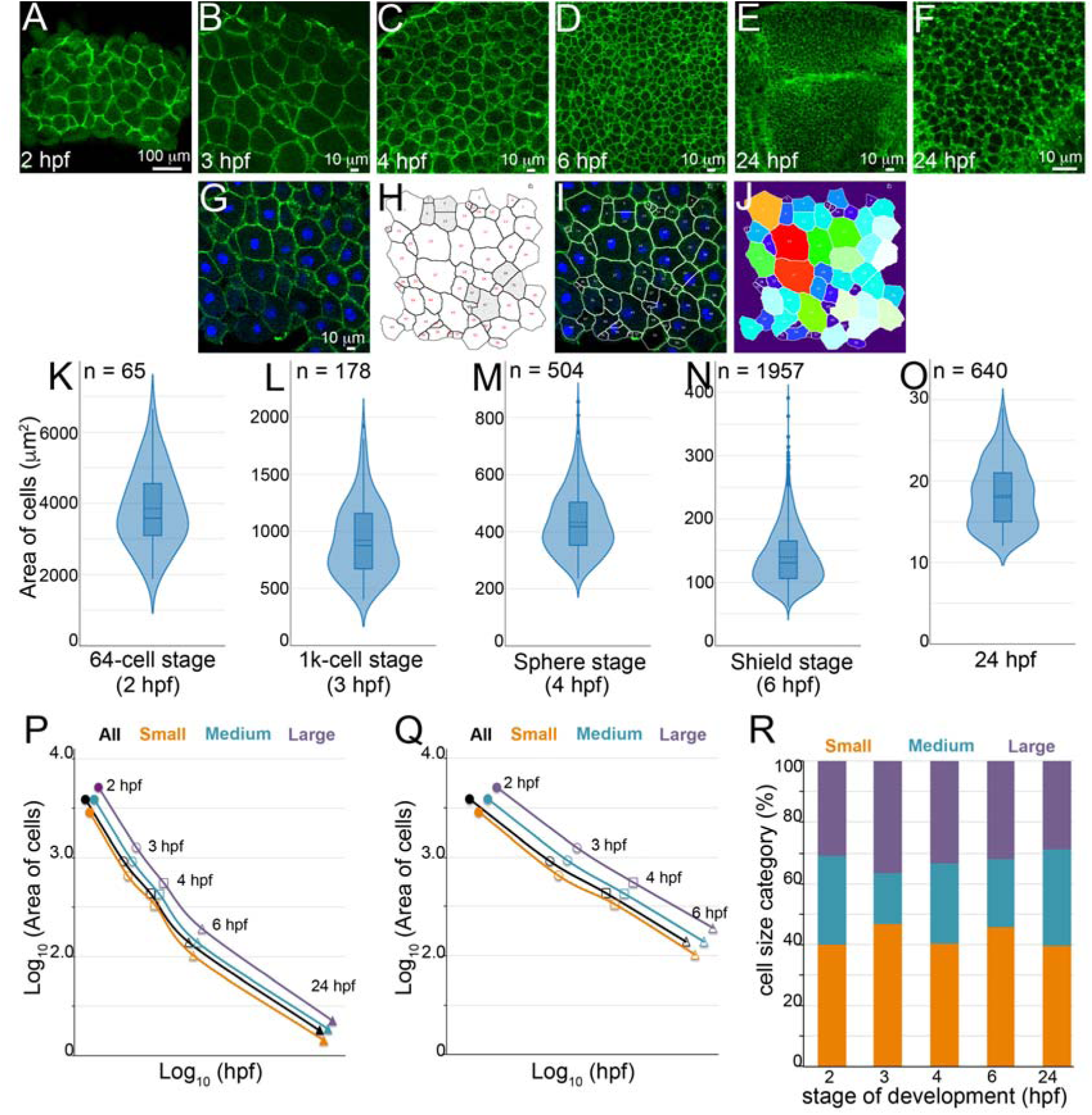
Quantification of cell size reduction during early development in diploid zebrafish embryos. β-catenin immunolabels of zebrafish blastoderms (A-F). Animal view of whole embryo at 2 hpf (A). One of the 4 quadrants imaged at 1k-cell (3 hpf, B), sphere stage (4 hpf, C) and shield stage (6 hpf, D) hpf. Left side of panel E is the mid-hindbrain boundary at 24 hpf and a higher magnification of top side of E (F). Quantification methodology to obtain cross-sectional areas of cells (G-J). Cell area distributions, dotted horizontal line within box plots indicate mean areas, n indicates total number of correctly segmented cells that were used for area calculations from multiple embryos (K-O). Cell size reduction plotted as log-log plots (P) and stages 2 to 6 hpf re-plotted to show linear reduction (Q), black line is the total cell population. Proportions of cell size categories during development (R).

**Figure 2:**
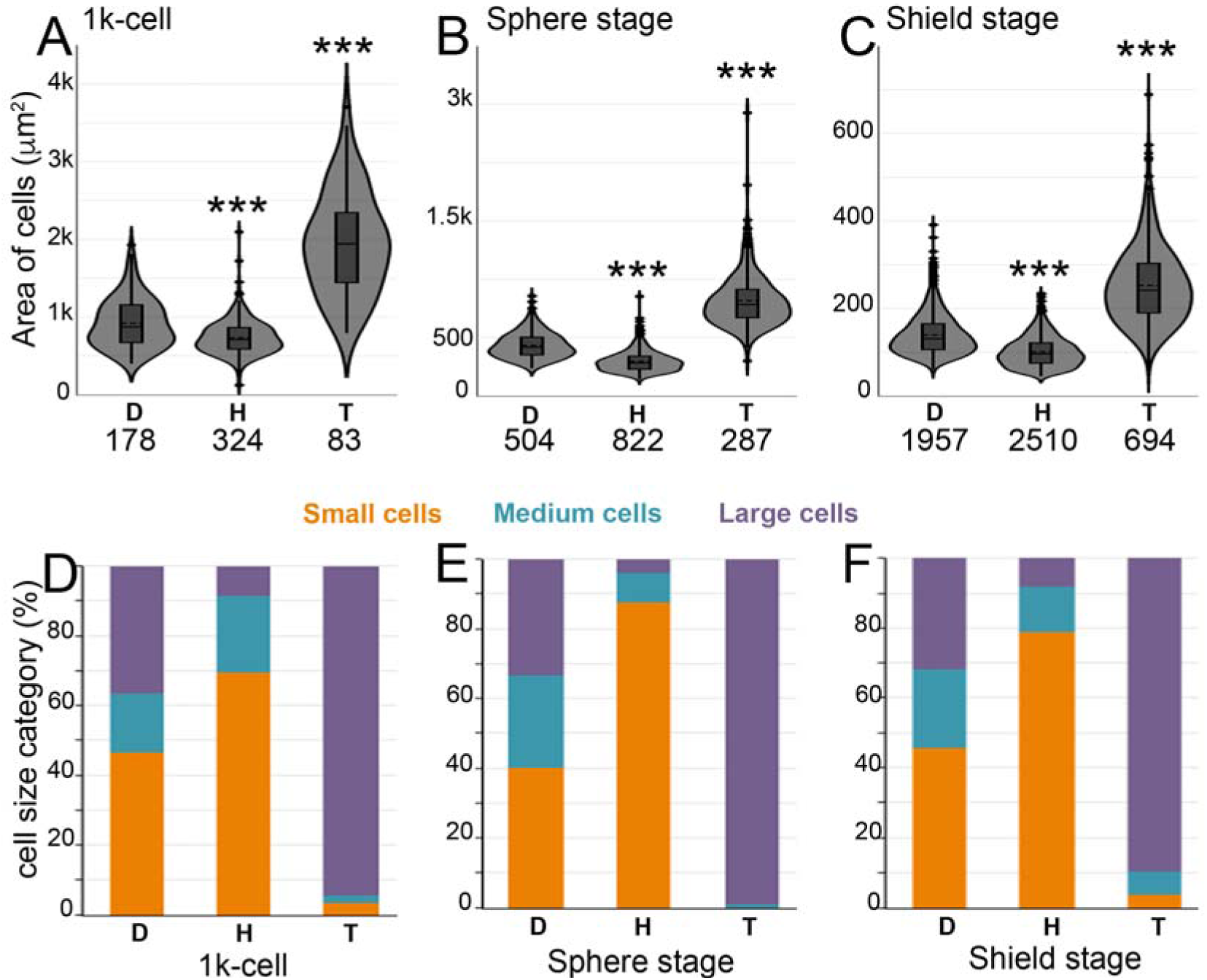
Ploidy alters the cell size norm during late blastula and gastrula stages of zebrafish development. Comparisons of cell area distributions in haploids and tetraploids to diploid normal values for a particular developmental stage, dotted horizontal line within box plots indicate mean area, n indicates total number of correctly segmented cells that were used for area calculations from multiple embryos (A-D). Asterisks denote statistical significance based on Mann Whitney (p ≤ 0.05). Proportions of cell size categories (F-J). (D) diploids, (H) haploids and (T) tetraploids.

**Figure 3:**
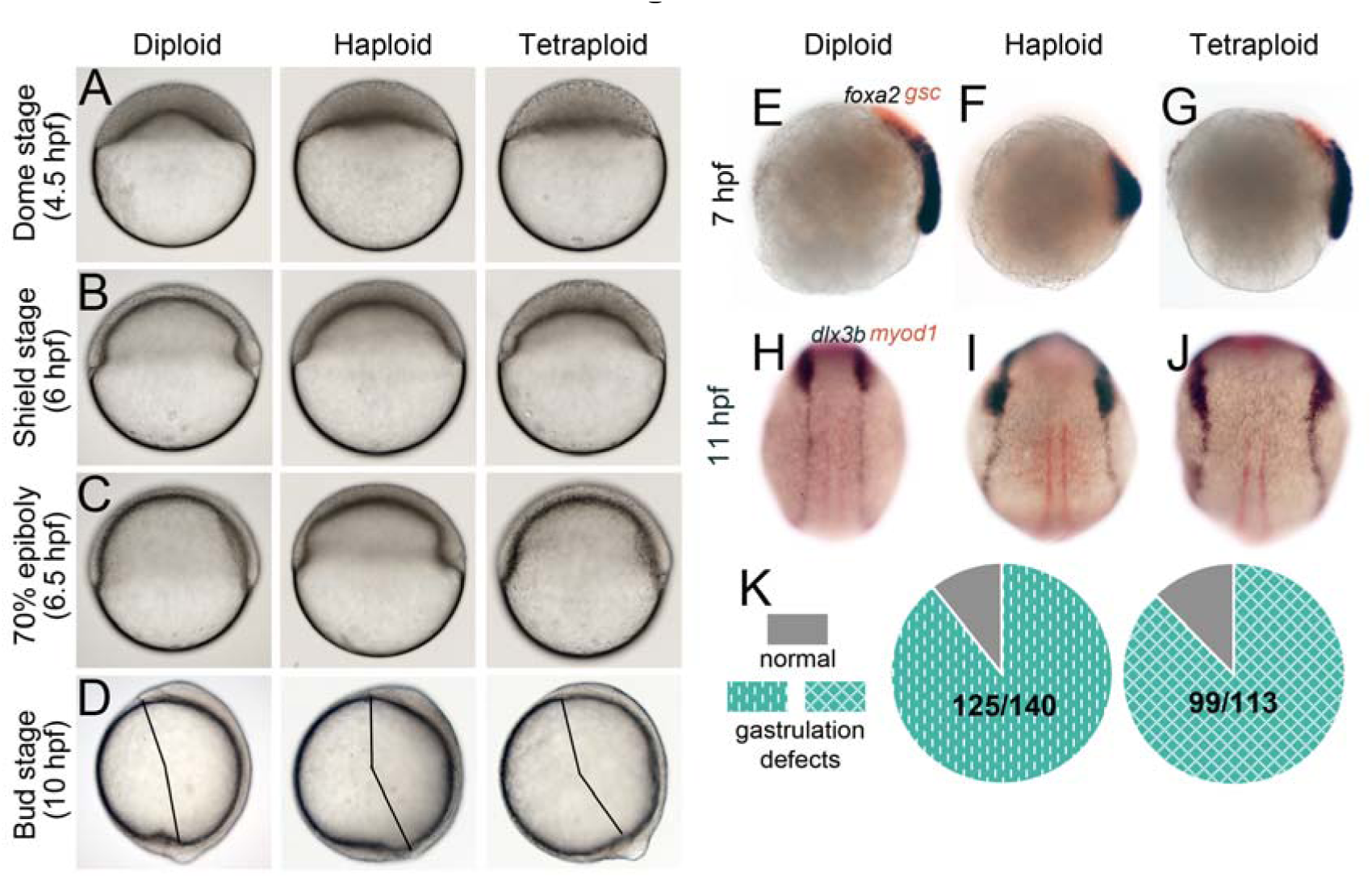
Embryos composed of smaller or larger than normal cells manifest epiboly and gastrulation defects. DIC images during epiboly and gastrulation (A-D) and gene expression analysis (E-K). For panels A-C, stages are noted according to diploid developmental stages and live appearance of haploids and tetraploids when diploids reached a particular developmental stage were recorded. Haploids and tetraploids initiate epiboly later than controls (A) and are delayed through gastrulation (B, C). For panels in D, the appearance of the tail-bud was taken as end of gastrulation and haploids and tetraploids were found to have a shorter body axis at this stage (D). RNA in situs for *foxa2* (blue) and *gsc* (red) at 7 hpf (E) and *dlx3b* (blue) and *myod1* (red) at 11 hpf (F). Quantification of gastrulation defects shown in panels F, I, G and J (K).

**Figure 4:**
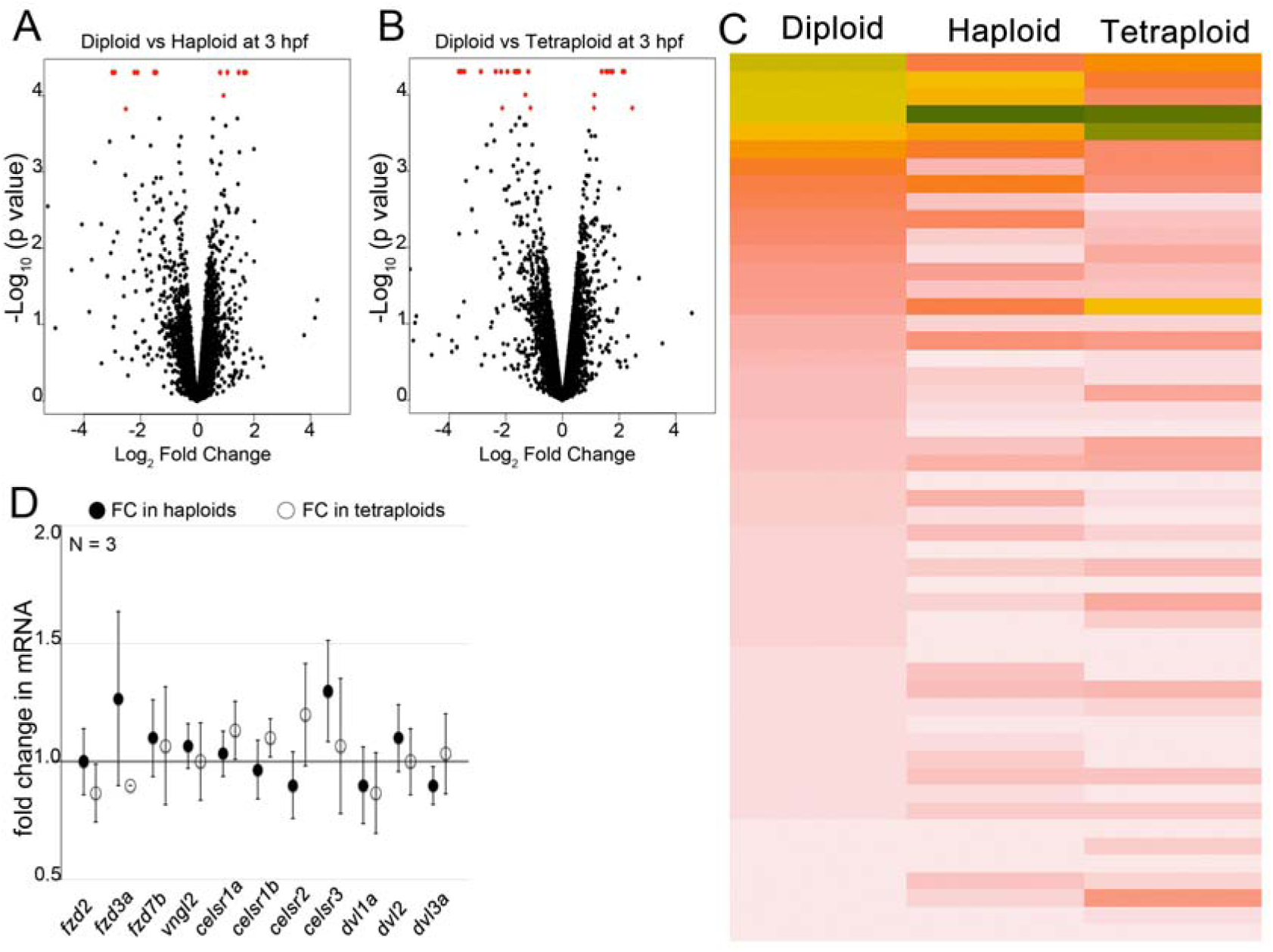
Transcriptional profiling and analysis of PCP pathway transcripts. Most genes are expressed normally in haploids and tetraploids just after ZGA at 3.5 hpf (black dots in A, B). Few genes are mis-regulated (red dots in A, B), which are genes differentially expressed with q values ≤ 0.05, the False Discovery rate corrected p-values. Heatmap of FPKMs of genes differentially expressed by Log_2_ fold change (C), arranged based on diploid expression level, decreasing from top to bottom. Quantitative RT-PCR at 8 hpf for PCP components revealed no significant change in transcript levels in comparison to diploids (D).

### Cell sizes in haploid and tetraploid zebrafish blastoderms deviate from the normal developmental stage-specific values

DNA content is known to affect cell sizes in varied contexts (Cavalier-Smith, 2005, Gregory, 2001, Lee et al., 2009, Otto, 2007). Experimentally generated haploid and tetraploid zebrafish embryos undergo normal rounds of early reductive divisions (Menon & Nair, 2018). 1k-cell, sphere and shield stages are reliably identifiable landmark stages during zebrafish development. These stages encompass developmental time points before initiation of cell migration (1k-cell), initiation of epiboly movements (sphere stage) and gastrulation movements (shield stage). Thus, potential cell size deviations from the norm at these stages could uncover the role of cell size in cell migration in developing embryos. Comparative analysis of cell sizes in haploids and tetraploids to diploid cell sizes at 1k-cell, sphere and shield stages revealed that cell sizes in haploid and tetraploid embryos were smaller and larger, respectively (Fig. 2A-C, Supplementary Fig. 1B, C, E, F, Supplementary Fig. 2B, C). During the first 6 hours of development, the average cell area in haploid embryos decreased by ∼36 fold and that in tetraploids decreased by ∼41 fold, with a decrease of ∼7-8 fold occurring between 1k-cell and shield stages (Fig. 2A-C). Binning cell size values from haploid and tetraploid embryos according to normal size values from diploid embryos showed that the small and large category of cells significantly deviated from the cell size norms across developmental stages (Supplementary Fig. 3A-D). Analysis of the proportions of cell size categories in haploids and tetraploid embryos revealed an over-representation of small cells in haploids and large cells in tetraploids, respectively (Fig. 2D-H). Regression analysis of reduction in cell area using log-log plots showed that cell sizes continued to decrease linearly suggesting power law reduction in cell size with development in haploids and tetraploids as well (Supplementary Fig. 4A-I).

### Haploid and tetraploid embryos develop epiboly and dorsal convergence defects

Reductive cell divisions are followed by collective cell migration during gastrulation, which rearranges cells in space over time to transform a collection of cells into an embryo. Collective cell behaviors are influenced by cell sizes in several different contexts, particularly during migration (Leal-Egana, Letort et al., 2017, Matsiaka, Penington et al., 2018). Cell size deviations in haploids and tetraploids occurred before gastrulation, allowing us to study the consequences of deviation from cell size norms on gastrulation. In diploid embryos, analysis of embryonic development in live embryos after sphere stages showed that diploid embryos underwent well-characterized collective cell behaviors such as migration over the yolk ball from the sphere stage (epiboly), followed by involution at the margin and anterior migration at the shield stage (initiation of gastrulation), and subsequent dorsal convergence to thicken and elongate the body axis on one side by the bud stage (end of gastrulation) (Fig. 3A-D). In haploids and tetraploids, initiation and progression of epiboly, formation of the dorsal shield and anterior migration were delayed (Fig. 3A-C). By the end of gastrulation, haploid and tetraploid embryos had a comparatively shorter body axis (Fig. 3D), which was also evident at 24 hpf (Supplementary Fig. 5A, C, E).

**Figure 5:**
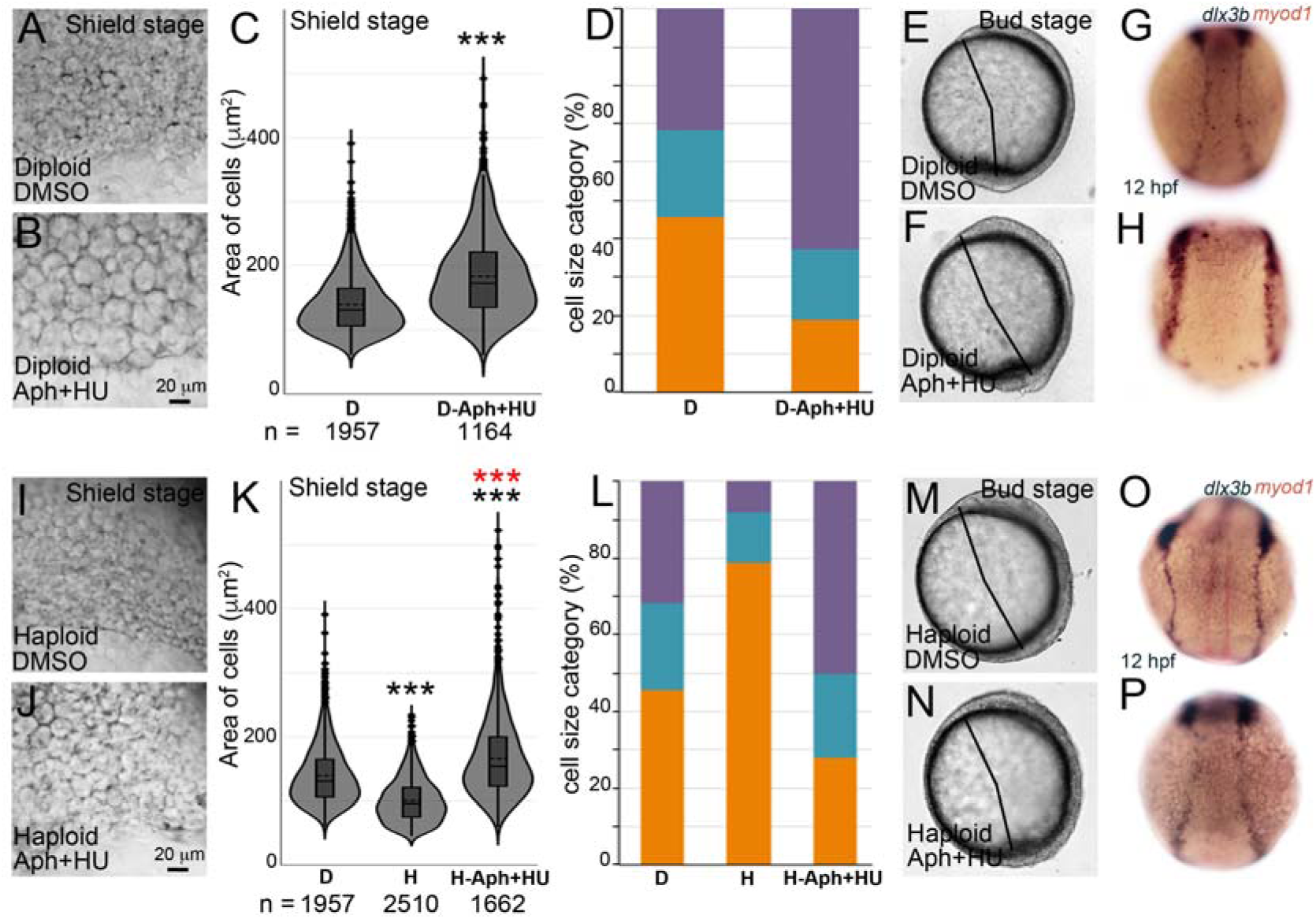
Developmental defects due alteration of stage-specific cell size norm can be recapitulated and rescued by transient pharmacological treatment. Diploids treated with 4% DMSO (A, E, G) or Aph+HU (B, F, H). At shield stages, cell sizes in diploid Aph+HU (B) are larger than in DMSO treated embryos (A). Cell area distributions in diploids and diploid Aph+HU at shield stages, dotted horizontal line within box plots indicate mean area, n indicates total number of correctly segmented cells that were used for area calculations from multiple embryos (C). Asterisks denote statistical significance based on Mann Whitney (p ≤ 0.05). Proportions of cell size categories at shield stsge (D). Diploid Aph+HU embryos have shorter body axis at end of gastrulation (FvsE) and ectoderm cells (blue in H, n=21/45) are further apart (G, n=29/29). Haploid embryos treated with 4% DMSO (I, M, O) or Aph+HU (J, N, P). At shield stages, cell sizes in haploid Aph+HU (J) are larger than in DMSO treated haploids (I). Cell area distributions in diploids, haploids and haploid Aph+HU at shield stages, dotted horizontal line within box plots indicate mean area, n indicates total number of correctly segmented cells that were used for area calculations from multiple embryos (K). Asterisks denote statistical significance based on Mann Whitney (p ≤ 0.05). Black and red asterisks indicate statistical significance in comparison to diploids and haploids, respectively. Proportions of cell size categories at shield stage (L). Bilaterally placed ectoderm cells (blue in P, n=21/27) are closer (O, n=16/26), indicating rescue of dorsal convergence.

Whole embryo RNA in situ hybridization to check for defects in anterior migration and dorsal convergence confirmed aberrant cell migration during gastrulation. At 7 hpf, the prechordal plate mesoderm cells at the anterior express *gsc*, and *foxa2* marks the posteriorly juxtaposed axial mesoderm (Fig. 3E). In haploids, the domains of *gsc* and *foxa2* expression remained intermingled and the *foxa2* expressing axial mesoderm cells were found in a shorter domain, indicating a failure in anterior migration of prechordal plate cells and elongation of axial mesoderm (Fig. 3F). In tetraploids, though there were no overt defects in *gsc* and *foxa2* expression domains, in general *gsc* expression seemed dampened (Fig. 3G). After completion of gastrulation at 10 hpf, the extent of dorsal convergence of cells can be assessed by the distance between the bilateral rows of ectodermal cells marked by *dlx3b*, across the dorsal midline (Fig. 3H). In both haploids and tetraploids, the rows of ectoderm cells were located further away from the midline, indicating a delay in or failure to converge dorsally (Fig. 3I, J). Thus, haploid and tetraploid embryos began undergoing delayed epiboly, and displayed a range of cell migration defects during gastrulation (Fig. 3K).

### Early transcriptomes of haploids and tetraploids and PCP pathway transcripts remain largely unperturbed

The experimental system used in this study to uncover the relevance of cell size in normal embryogenesis was alteration of ploidy in zebrafish embryos. This experimental paradigm was chosen primarily because there are no zebrafish mutants with smaller or larger cells during 1k-cell to shield stages that would allow us to study the consequence of altering cell sizes before and during gastrulation on overall embryonic development. DNA content has been shown to affect timing of zygotic genome activation (ZGA), which occurs at 2.5 hpf in zebrafish (Kane & Kimmel, 1993, Newport & Kirschner, 1982a, Newport & Kirschner, 1982b). To prove that gene expression levels in haploids and tetraploids remained comparable to diploids, we performed whole transcriptome analysis at ∼3.5 hpf. Whole transcriptome comparisons showed that at ∼3.5 hpf, majority of genes were not mis-expressed in response to ploidy alteration (Fig. 4A, B). 21 genes in haploids and 33 genes in tetraploids were mis-expressed (Fig. 4C and Supplementary Table 1). Interestingly, only 6 genes are mis-regulated in common between haploids and tetraploids, yet both kinds of embryos manifest similar defects later in development (discussed in later sections), suggesting that the developmental defects do not arise from perturbed ploidy *per se*. Dorsal convergence, which is compromised in both haploids and tetraploids, is most pronounced during late gastrula stages (Kimmel et al., 1995). A molecular pathway known to drive collective cell migration in diverse cellular and evolutionary contexts, including dorsal convergence is the Planar Cell Polarity (PCP) pathway (Davey & Moens, 2017, Heisenberg, Tada et al., 2000, Topczewski, Sepich et al., 2001). Quantitative real time PCR showed that transcript levels of 11 genes representing diverse components of the PCP signaling pathway remained unaltered in haploids and tetraploids at 8 hpf (Fig. 4D).

### Drug-induced cell size increase in diploid blastoderm recapitulates gastrulation defects

Gene Ontology analysis of the 54 mis-expressed genes in haploids and tetraploids showed that these genes controlled a diverse range of biological functions (Supplementary Table 1). However, an unbiased approach to test whether deviation from cell size norms during early development could result in gastrulation defects would be to recreate the cell size deviation phenomenon in diploids. A combinatorial treatment of Aphidicolin and Hydroxyurea (Aph+HU) has been used to effectively increase cell sizes in the zebrafish notochord (Liu, Sepich et al., 2017). To preclude the possibility that exposure to Aph+HU at early developmental stages was inducing DNA damage, we assayed for γH2AX foci in the nuclei after Aph+HU treatments. Diploid zebrafish embryos treated with Aph+HU from 1k-cell to shield stages showed negligible toxicity and DNA damage (Supplementary Fig. 6, 7). Average cell areas in Diploid Aph+HU embryos were larger than normal at shield stages, with large cells representing majority of the sample population (Fig. 5A-D, Supplementary Fig. 3E). By bud stages, diploid Aph+HU embryos had shorter body axis, which resembled the phenotypes seen in haploids and tetraploids (Fig. 5E, F, compare to Fig. 3D). Analysis of the extent of dorsal convergence at 12 hpf showed that, the *dlx3b* expressing ectoderm was found further away from the midline, a phenotype that also resembled the *dlx3b* expression in haploids and tetraploids (Fig. 5H, compare to Fig. 3I, J). Thus, in a ploidy-independent experimental paradigm, increasing cell sizes in diploid embryos between 1k-cell to shield stages also resulted in gastrulation defects similar to those seen in haploid and tetraploid embryos.

**Figure 6:**
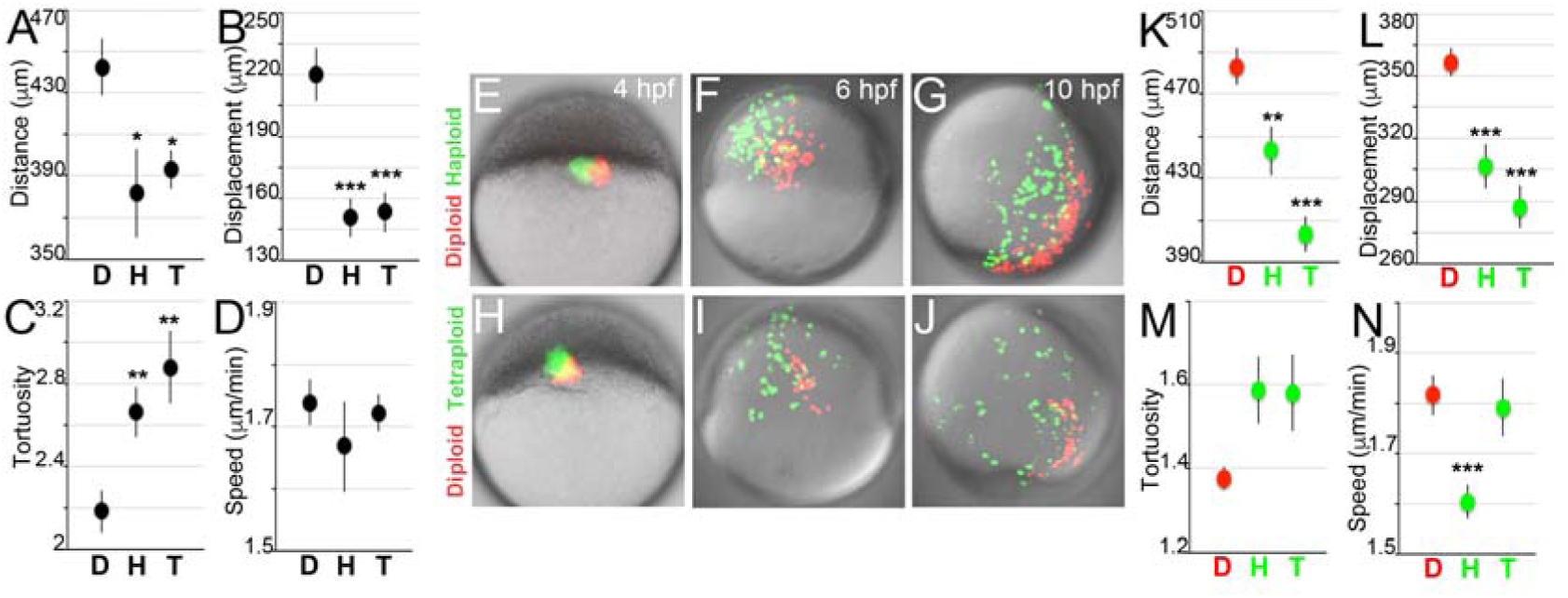
Smaller or larger than normal cells migrate sub-optimally during gastrulation. Tracking of cell migration in clonally labeled embryos (A-D) and in chimeric embryos (E-N). Graphs show total distance travelled (A), net displacement during migration (B), tortuosity (C) and average speed (D) of clonally labeled cells in diploids (D), haploids (H) and tetraploids (T) from shield to bud stages of gastrulation. Diploid (red) cells co-transplanted with haploid or tetraploid (green) cells into diploid hosts at sphere stage (4hpf, E, H) and imaged through gastrulation (F, G, I, J). Graphs show total distance travelled (K), net displacement during gastrulation (L) tortuosity (M) and average speed (N) of diploid (D, red), haploid (H, green) and tetraploid (T, green) cells. Asterisks denote statistical significance based on Mann Whitney (p ≤ 0.05).

**Figure 7:**
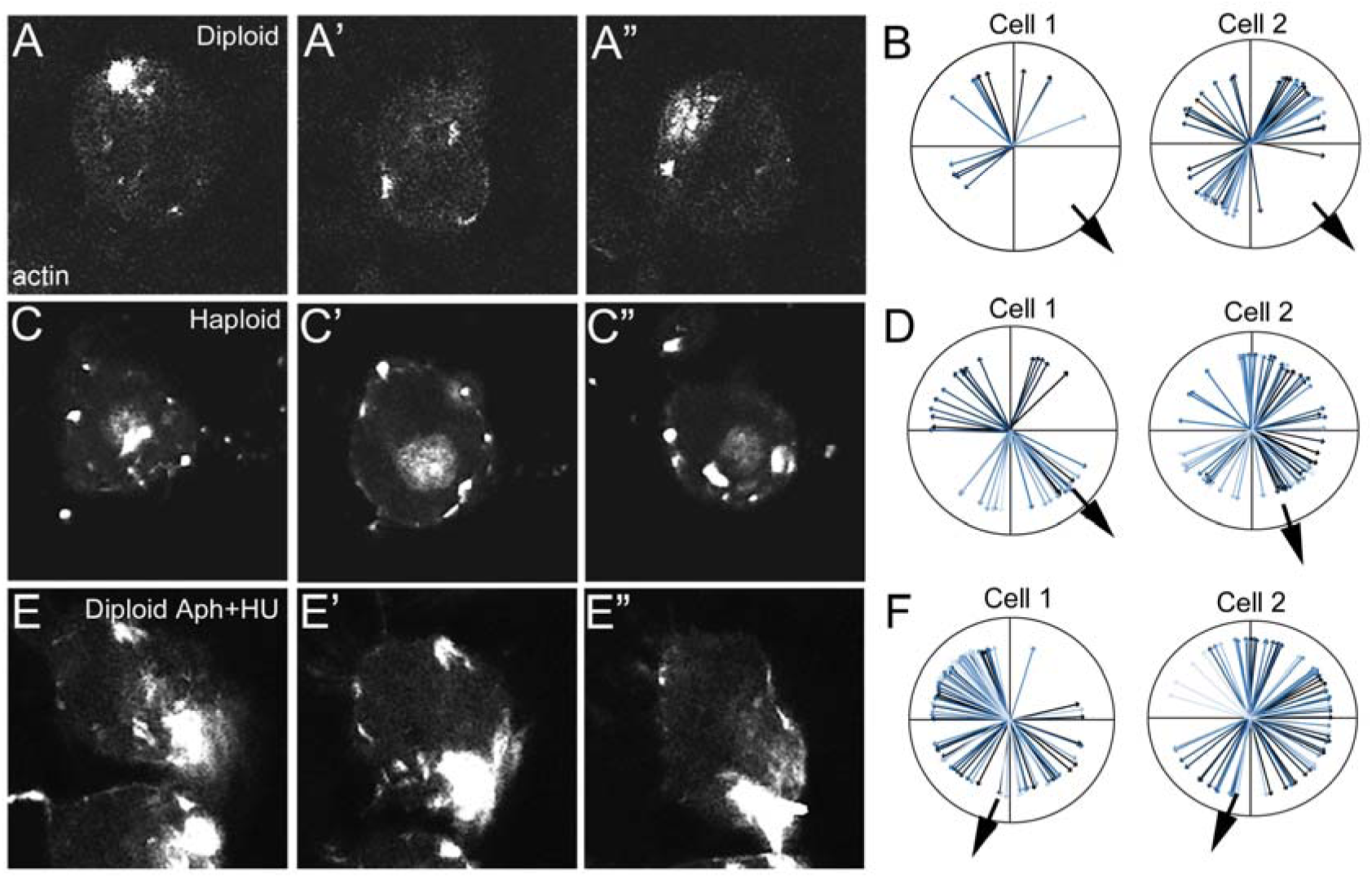
Smaller or larger than normal cells mis-localize actin during migration. Dynamic actin localization changes in migrating cells (A-F). Panels A, C and E are higher magnification still frames of one cell during migration from Supplementary Movies 6, 9 and 12, respectively. Actin localization in migrating cells over time shown as lines drawn at angles from the center of a cell, black arrowhead indicates the direction of migration in the movie.

**Figure 8:**
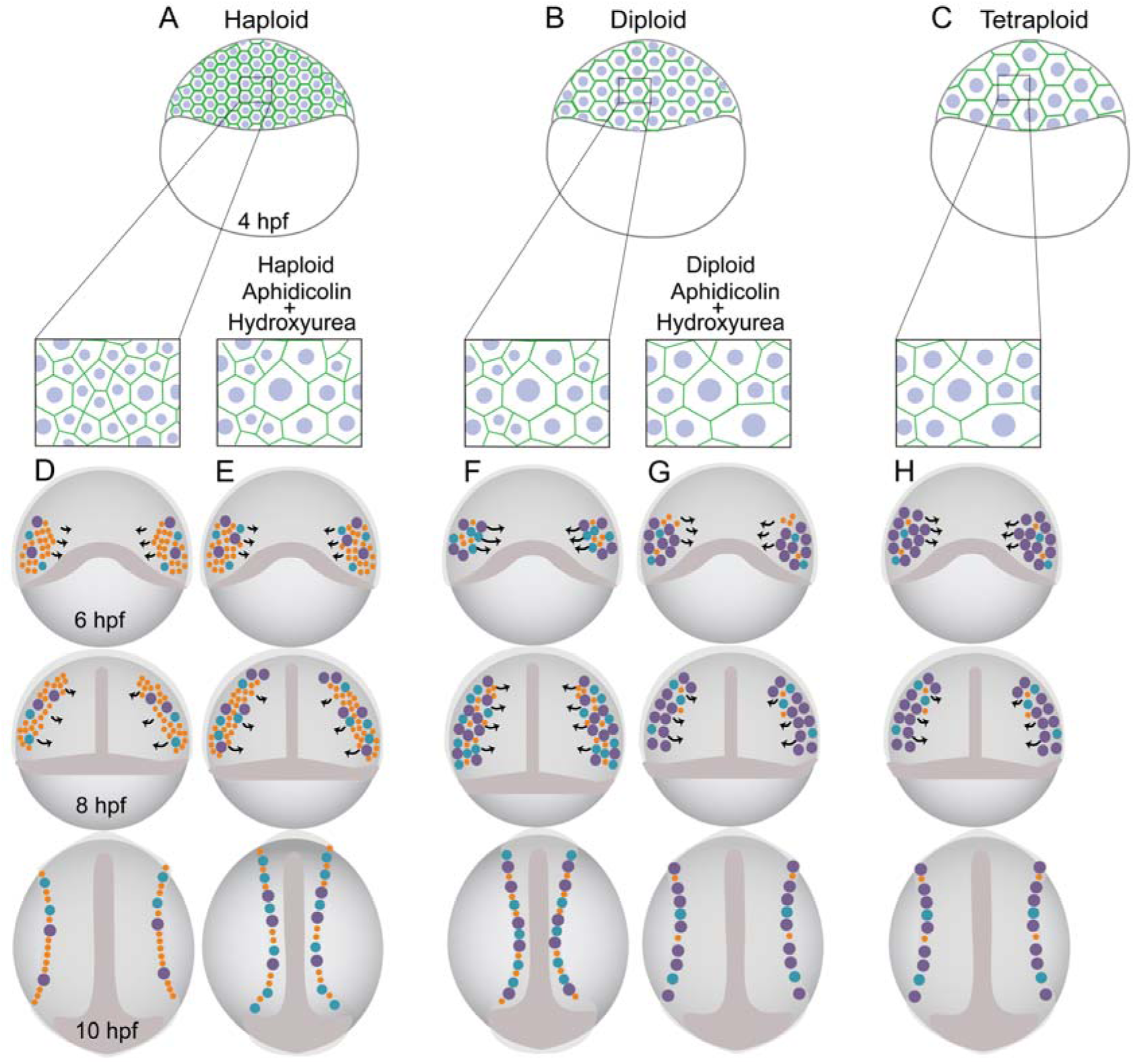
A landscape of optimal cell sizes enables efficient collective cell migration for normal development. Haploid (A), diploid (B) and tetraploid (C) embryos with smaller cells in haploids (box in A), normal cell sizes in diploids (box in B) and larger cells in tetraploids (box in C). Diploid embryos (F) made of small (orange), medium (blue) and large (purple) cells in normal proportions, have stage-specific normal size ranges and gastrulate normally. Haploid and Tetraploid embryos deviate from the stage-specific cell size norm, have more small (D, orange) or large (H, purple) cells, and manifest epiboly and dorsal convergence defects during gastrulation (D, H). Aph+HU increases cell size (rectangular panels above E and G). In diploids, this cell size increase recapitulates in diploids gastrulation defects (G), while in haploids, this partially rescues gastrulation defects (E).

### Drug-induced cell size increase in haploid blastoderms rescues gastrulation defects

If cell size deviations from the norm during 1k-cell to shield stages does indeed cause gastrulation defects, then restoring cell sizes to the norm may rescue the developmental defects. Cell size increase upon Aph+HU treatment allowed us to test this experimentally in haploid embryos, wherein cell sizes are smaller than the norm between 1k-cell and shield stages. Aph+HU treatment of haploid embryos from 1k-cell to shield stages (3 hour exposure) as was done for diploid embryos resulted in cell extrusions during gastrulation indicating that haploid embryos were fragile. A shorter Aph+HU exposure regime from 1k-cell to dome stage (1.5 hour exposure) was therefore used for haploid embryos. Haploid embryos treated with Aph+HU from 1k-cell to dome stages showed an increase in cell sizes to diploid-like values by the shield stage (Fig. 5I-K, Supplementary Fig. 3F). Rescuing the cell areas in haploid Aph+HU to diploid-like values by shield stage also restored the proportions of cells within different size categories to diploid-like proportions (Fig. 5L). Importantly, gastrulation defects were also partially rescued in haploid Aph+HU embryos (Fig. 5M, N). Analysis of dorsal convergence in haploid Aph+HU embryos clearly showed that the ectoderm marked by *dlx3b* was closer to the midline indicating a rescue in dorsal convergence (Fig. 5O, P).

### Smaller or larger than normal cells migrate sub-optimally during gastrulation

To assay cell migration in embryos with smaller or larger size cells than the norm, fluorescently labeled cell clones were generated in diploid, haploid and tetraploid embryos and migration of individual cells were tracked during gastrulation (Supplementary Movies 1-3). Quantification of cell migration parameters revealed that cells in haploids and tetraploids covered less distance (total path length travelled) and were also less displaced (net change in position from beginning to end of imaging) in comparison to cells in diploid embryos (Fig. 6A, B). The observed deficiencies in distance covered and displacement could arise because cells move slower and/or took more tortuous paths during migration. In comparison to cells in diploid embryos, cells in haploids and tetraploids did indeed take significantly more tortuous paths during migration (Fig. 6C). The average speed of cells in tetraploids was comparable to cells in diploids, whereas the average speed of cells in haploids tended to be slightly slower (Fig. 6D).

An advantage of zebrafish as a model organism is the ease of generating chimeric embryos to assay experimental outcomes when cells are provided developmental milieus different from their native environments. We next tracked cell migration during gastrulation in chimeric diploid embryos in which small groups of differentially labeled diploid:haploid (red, green) or diploid:tetraploid (red, green) cells were co-transplanted. Cells were transplanted into diploid host embryos at sphere stage, just prior to initiation of epiboly, and tracked live by timelapse imaging till the end of gastrulation. Though the initial positions of the normal-small (diploid:haploid) and normal-large (diploid:tetraploid) cells were the same at sphere stages (Fig. 6E, H), during gastrulation smaller and larger cells were found farther laterally in the embryo in comparison to co-transplanted normal size cells (Fig. 6F, I and Supplementary Movies 4, 5). By the end of gastrulation, smaller and larger cells were still displaced farther laterally and were consistently found further away from the midline in comparison to the co-transplanted normal cells (Fig. 6G, J and Supplementary Movies 4, 5). Quantification of cell migration parameters of co-transplanted normal-small and normal-large cells revealed that small (haploid) and large (tetraploid) cells traveled less distance (Fig. 6K), were displaced less (Fig. 6L) and took significantly more tortuous paths (Fig. 6M), in comparison to normal (diploid) cells. The average speed of large cells was comparable to normal cells whereas the average speed of small cells was significantly slower than normal size cells (Fig. 6N).

### Smaller or larger than normal cells show altered actin dynamics

Smaller or larger than normal cells may migrate aberrantly due to altered actin dynamics during collective cell migration. Therefore, actin dynamics during gastrulation was assayed by generating mosaic diploid embryos in which diploid (normal), haploid (small) or diploid Aph+HU (large) cells were transplanted at sphere stages and imaged live during gastrulation (Supplementary Movies 6-14). The transplanted cells were fluorescently labeled for cell membrane and actin. In normal (diploid) cells, an actin patch formed at the back of the cell, which rearranged into smaller dynamic membrane punctae and then reassembled as a patch at the back of the cell. To better understand the dynamic actin localizations, azimuthal distribution of actin patches within single cells was analyzed with respect to the direction of migration. This analysis revealed that actin localized to 3 of the 4 quadrants in a cell, leaving one quadrant, in the direction of migration, largely free of actin during migration (Fig. 7A-B, Supplementary Movies 6-8). Similar analysis of the positions of actin in migrating small cells (haploid) revealed several larger actin patches in all four quadrants of the cell, which moved randomly without disassembly during migration (Fig. 7C-D, Supplementary Movies 9-11). In large cells (diploid Aph+HU), a very large patch of actin formed at the front of the cell and remained at the front without disassembling during migration in addition to the actin punctae that localized to all four quadrants during migration (Fig. 7E-F, Supplementary Movies 12-14).

## DISCUSSION

A quantitative analysis of the evolution of progressive reduction in cell sizes during development revealed that early zebrafish embryos are composed of heterogeneously sized cells, which exist in certain proportions. We do not know how these proportions are set in the developing embryo and the relevance of the proportions of cell size categories to normal development. The quantifications of cell areas from a sample cell population within the deep cell layer also revealed that cell size reduction leading up to gastrulation is linear, following power law behaviour. The cellular and molecular mechanisms that regulate cell size dynamics during early embryonic development are not known in any metazoan. Because alternate methodologies to alter cell sizes during the early reductive cell division phase of development are unavailable we used haploid and tetraploid zebrafish embryos together with pharmacological perturbations to alter cell sizes. Altered ploidy zebrafish embryos are advantageous for this study because unlike mouse haploid and tetraploid embryos, which are early embryonic lethal, zebrafish haploids and tetraploids are late larval lethal (Eakin, Hadjantonakis et al., 2005, Henery et al., 1992, Kaufman & Gardner, 1974, Koizumi & Fukuta, 1995, Modlinski, 1975). As seen in multiple cell types in both plants and animals (Henery et al., 1992, Orr-Weaver, 2015), we found that cell size in zebrafish haploids and tetraploids correlated with the ploidy. Such deviation in cell size was uniquely advantageous to this study as it allowed assessment of the consequence of deviating from a cell size norm early in development on subsequent embryogenesis in an endogenously diploid vertebrate. In haploids and tetraploids absolute cell sizes as well as proportions of cells across size categories deviated from that in diploids at comparable developmental stages. Haploid and tetraploid zebrafish embryos were delayed in development during late blastula and early gastrula stages. Morphological defects exacerbated with time and culminated in dorsal convergence defects by the end of gastrulation. This study does not distinguish the possibility of developmental defects due to cell size deviations versus due to perturbations in the proportions of size categories. We interpret the developmental defects as arising from a combination of the deviations from the normal cell size and proportion norm for a developmental stage.

Comparative transcriptomics in haploids and tetraploids just after ZGA revealed that levels of majority of the transcripts remained comparable to diploids, suggesting no gross transcriptional perturbations. This is also true with respect to ploidy in yeast (Galitski, Saldanha et al., 1999). Interestingly, despite the fact that haploids and tetraploids manifest similar gastrulation defects, mis-regulated transcripts between the two conditions are largely unique. Therefore, independent transcriptional cascades may go awry in haploids and tetraploids and yet cause similar aberrant developmental outcomes. Alternatively, transcription independent mechanisms may regulate the evolution of cell size dynamics during early development. A complementary, ploidy-independent alteration of cell sizes in diploid embryos by transient pharmacological treatments support the interpretation that independent of gene expression changes, deviations in cell size from the norm during early stages can cause consistent developmental abnormalities. It would be quite unlikely that alteration in ploidy and pharmacological perturbations both result in cell size deviation from the norm and gastrulation defects due to general toxic or pleiotropic effects. Such an interpretation is strengthened by a phenotypic rescue of haploids wherein gastrulation defects were mitigated when cell sizes were restored to normal-like sizes at shield stages. It is again highly unlikely that two separate and independent sets of potential pleiotropic effects (arising from altered ploidy and pharmacological perturbations) would add up to rescue developmental defects in haploids.

The developmental defects we describe do not resemble those seen when apico-basal polarity or cell adhesion is perturbed in zebrafish embryos (Kane, McFarland et al., 2005, Sonawane, Carpio et al., 2005). Recent work in size reduced zebrafish blastoderms showed that signaling gradients dynamically scale to readjust somite sizes to scale with smaller embryo sizes (Ishimatsu, Hiscock et al., 2018). In *D*. *melanogaster*, morphogen gradients scale in response to cell size change to maintain a normal wing size (Day & Lawrence, 2000). In early zebrafish embryos, when cell sizes deviate from a developmental stage-specific norm, perhaps cells perceive signaling gradients in the blastoderm or early gastrula differently. However, regardless of sensing the deviation from a cell size norm, embryos are unable to compensate for the suboptimal early cell sizes to restore normal patterning.

Regulating cell size during early development when cell size is a constantly changing parameter versus regulating cell size during homeostasis where the goal is to maintain a certain cell size may require distinct mechanisms. It is possible that the dynamic nature of cell size change during early development uses an equally dynamic cytoskeletal interaction between astral microtubules and cortical actin to sense cell boundaries and adjust cell size dynamically, especially in the rapidly dividing zebrafish embryo. The spatial analysis of gene expression domains indicated compromised cell migration when developmental stage-specific cell sizes deviated from diploid-like values. Tracking cell migration during gastrulation in chimeric and mosaic embryos during gastrulation revealed that smaller or larger cells migrated sub-optimally in comparison to normal size cells and was likely to cause aberrant positioning of cells during gastrulation, resulting in phenotypes such as compromised dorsal convergence. Additionally, actin reorganization during migration was altered in cells that were smaller or larger than normal, suggesting that cell size deviation from the norm affects cell migration at least partially by altering actin dynamics. The mis-localization of actin into all four quadrants in smaller or larger cells may drive cells to take tortuous paths during migration, making the whole process of migration suboptimal. Taken together, live tracking of cell migration during gastrulation in zebrafish indicates that cell migration parameters are cell autonomous and may depend on the size of the cell.

It is intuitive that during early embryogenesis, a phase characterized by growing numbers of progressively smaller cells, optimum cell sizes may be important for normal development. However, in the absence of experimental efforts directed at altering cell sizes during the earliest phases of development, this hypothesis remains an intuitive conjecture at best. Reductive cell division after fertilization is an evolutionarily conserved phenomenon in metazoans. Our study shows that zebrafish embryos made of smaller or larger than normal cells develop abnormally, likely due to cell autonomous and possibly size-dependent effects on migratory properties of cells, which affects gastrulation. In developing embryos, early reductive cell divisions potentially allow dynamic, developmental stage-specific cell size norms to emerge. Perhaps such a landscape of optimal cell sizes enables efficient collective cell migration to correctly position cells in space and time to shape the amorphous ball of cells in a blastoderm into a patterned embryo at the end of gastrulation.

## MATERIALS AND METHODS

### Zebrafish

Adult *Danio rerio* of the standard laboratory strains Tubingen and AB were reared at 28.5°C under 14 hour light and 10 hour dark conditions. To synchronize developmental stages, embryos were obtained by in vitro fertilizations (IVF) for all experiments and raised at 28.5°C. Diploid, gynogenetic haploid and tetraploid embryos were generated as described in (Menon & Nair, 2018, Menon & Nair, 2019).

### Validation of ploidy

24 hpf larvae were dechorionated and arrested at metaphase in 4mg/mL Colchicine (Sigma C9754) at 28°C for 90 min in the dark. The larvae were treated with 1.1% sodium citrate monobasic and yolks were punctured in a span of 8 mins at room temperature (RT) followed by incubation on ice for an additional 8 mins. Deyolked embryos were fixed overnight in 3:1 Methanol:Acetic acid at 4°C, transferred into 50% Acetic acid and minced thoroughly with forceps. A single cell suspension was generated by repeated trituration using 50 μl glass capillary pipettes and 2-3 drops were dropped from a height onto a glass slide pre-warmed to 65°C. The slides were incubated at 65°C for 60 mins and stained with DAPI (Invitrogen P36974) overnight at RT. Metaphase chromosomes were imaged on Zeiss Axio-Imager M2 and counted using Image J.

### Developmental Staging of embryos

To stage-match embryos for comparative analysis IVF embryos at 1k-stage (3 hpf), high stage (3.5 hpf), dome stage (4.5 hpf), sphere stage (4 hpf) and shield stage (6 hpf), were staged according to the characteristic morphology of the embryo at these landmark stages. For 11 and 12 hpf fixations, the somite number was used as the criterion for stage matching embryos. 2 and 24 hpf embryos were staged by absolute time after fertilization.

### Immunofluorescence and imaging

Embryos were fixed using 4% paraformaldehyde and immunolabelled with rabbit anti-β-catenin (Sigma C2206, 1:1000) or rabbit anti-γH2AX (Cell Signaling Technology 2577S, 1:1500) and donkey anti-rabbit Alexa 488 (Invitrogen A21202, 1:100) (Pelegri, Knaut et al., 1999). DNA was stained using DAPI. Semi-flat mounts of the blastoderms were prepared in 70% glycerol containing DABCO (Sigma D27802) and imaged on an Olympus FV1200. Images were processed and analyzed using ImageJ and figure panels were assembled using Adobe Photoshop. For 1k-cell, sphere and shield stage embryos, all four quadrants were imaged for cell size quantifications to cover majority of the blastoderm.

### Cell size quantifications

To quantify the cell area of the deep layer cells, a single z-slice from the middle 2/3^rd^ of a β-catenin and DAPI labeled confocal z-stack was chosen from all images. This z-slice was segmented using the macro http://imagej.1557.x6.nabble.com/Index-of-hexagonality-SOLVED-td5013442.html. The segmented confocal z-slice was an image in which each cell was outlined and numbered. These outlined and numbered cells were overlaid on the original confocal image to verify the segmentation. Aberrantly segmented cells were eliminated or manually outlined and numbered. To obtain maximal cross-sectional area of a cell, areas were only calculated from cells in which the nucleus was clearly visible. The correctly segmented image was used in the Extended Particle Analyzer and Shape Descriptor Map plugins of the Biovoxxel Toolbox in ImageJ which generated area heat maps and an excel sheet with the cross-sectional area of each numbered cell. 3 to 4 embryos were analyzed for 2 and 24 hpf stages. All four quadrants from 3 embryos each were analyzed for 1k-cell, sphere and shield stage embryos. Area calculations were done using Microsoft Excel. Since the data was not normally distributed, statistical significances were calculated using Mann Whitney and Kruskal Wallis tests in Graphpad Prism. Violin plots were made using Chart Studio at https://plot.ly

### Whole mount RNA in situ hybridization

Embryos fixed in 4% paraformaldehyde were processed for whole mount in situ hybridization (Thisse, Thisse et al., 1993). Digoxigenin or Fluorescein labeled anti-sense RNAs for the genes described were detected using anti-digoxigenin (Roche 16646820) and anti-fluorescein antibodies (Roche 17110000), using NBT (Roche 11383213001) and BCIP substrate (Roche 11383221001) or Sigma Fast (Sigma F4648). Stained embryos were post-fixed in 4% PFA and mounted in 100% glycerol for imaging on Zeiss AxioImager M2. Images were processed and assembled using ImageJ and Adobe Photoshop.

### Transcriptome analysis

50 pairs of adults of the AB strain were used exclusively to generate diploid, haploid and tetraploid embryos for transcriptome sequencing. For one biological replicate, embryos were obtained from ∼15 pair matings and diploid, haploid and tetraploid embryos were generated from the same clutch of embryos as described in (Menon & Nair, 2018). At high stage, total RNA was extracted from ∼120 embryos of each ploidy by Trizol method (Ambion 15596018). RNA quality was assayed on a Bioanalyzer (Agilent 2100). The RNA set in which RNA from diploid, haploid and tetraploid embryos generated in the same experiment had RIN values >/= 8-9 was sent for whole transcriptome sequencing. The entire experiment of generating all three ploidies and RNA extraction was repeated two more times on different days to obtain a biological replicate of three RNA samples for each ploidy. Ploidy was verified by metaphase chromosome spreads at 24 hpf for each sample that was used for transcriptome sequencing. Strand specific library was generated and 15-20 million 100 bp paired-end Illumina reads were obtained on HiSeq 4000. The Illumina reads were processed and analyzed using HISAT2 and Cuffdiff at the Galaxy open source platform (https://usegalaxy.org/) to obtain differentially expressed transcripts from the three biological replicates of transcriptomes of each ploidy. R software was used to generate the Volcano plots and heatmap.

### Quantitative RT-PCR

Diploid, haploid and tetraploid embryos were generated from the same IVF clutch as was done for the transcriptome sequencing. Total RNA was extracted from ∼40 diploid, haploid, and tetraploid embryos using Trizol. 1μg of the RNA converted into cDNA using the SSIII RT First Strand cDNA synthesis kit (Invitrogen 18080051). qRT-PCR was performed using Kapa Sybr (Kapa Biosystems KK4601) reaction mix on a Roche LightCycle 96. The q-RTPCR primers used were: *fz2*-F-GAGTTTGCTTTGTGGGCCTG and *fz2*-R-GCAAGCAGAAATG AGGTGCC, *fz3a-*F-A GAGCGACTGCTACAAGCTC and *fz3a-*R-GCTCGAGGGTACGACTCTTC, *fz6-*F-AAGT ACCTGATGACGCTGGC and *fz6-*R-CGCTGACCGCATCTTTCTTG, *fz7b-*F-AGGATAG TTTGTGCCTCGCC and *fz7b-*R-TTCCCGTACCGCCATCATTG, *vangl2*-F-AGCCGCTT CTACAATGTGGG and *vangl2*-R-TGGGAAGGTTGAGTAAAGCAGG, *celsr1a*-F-TCC ACGAGAACATCTGAACGG and *celsr1a*-R-TCCTGCTGGGTCAGATTGATG, *celsr1b*-F-ACCTGGATAACAACAGGCCG and *celsr1b*-R-TGTTCTCCAGACGAACGGTG, *celsr2*-F-CTTTCATCGAGGGCGGAGTC and *celsr2*-R-ACGAGTCACAGTCGTCAACC, *celsr3*-F-GCGTCTCCGAAGTGTCACC and *celsr3*-R-GATCGCTCCAGCTGCTTTTG, *dvl1a*-F-ACGATGATGCCAGCAGACTC and *dvl1a*-R-TTGTGGTCTTTCGGTCTCCG, *dvl2*-F-TCCTTACTCCACCCAGCCTC and *dvl2*-R-AACGCTTTTAGGGCCCCTTC, *dvl3a-*F-AGTTGCTTTGCTTTGCCGAG and *dvl3a-*R-GGATGGACTCATGCCGTAGG, *dvl3b*-F-GGGATCCAAACCCTCGTAGC and *dvl3b*-R-GGCTCATACCGTACACTGGG, *tbp*-F-ACCTCCTTTCGCTCAAGGTC and *tbp*-R-CCAGG AGGGACAAGCTGTTG. The fold-change of test transcripts was analyzed with respect to *tbp* transcript levels. The data was analyzed and graphically represented using Microsoft Excel.

### Pharmacological treatments

Standardization experiments for Aphidocolin (Sigma A0781) and Hydroxyurea (Sigma H8627) (Aph+HU) treatment of zebrafish embryos during blastoderm stages is shown in Supplementary Fig. 6. Before addition of the drugs, embryos were transferred into 0.3% Danieau medium (17mM NaCl, 0.21mM KCl, 0.12mM MgSO_4_.7H_2_O, 0.18mM Ca(NO_3_)_2_.4H_2_O and 1.5mM HEPES; pH 7.6) and dechorionated using Pronase (Sigma P6911). At 1k-cell stage, Danieau medium was replaced with ∼10 ml of 50 μM Aphidicolin and 5 mM Hydroxyurea solution prepared in 4% DMSO. This concentration of Aph+HU was used for all experiments after standardization. Control embryos were treated with only 4% DMSO. The drug solution or DMSO was discarded at shield stage for diploids or at 4.5 hpf for haploids and for diploid embryos used as donors in the actin timelapses. Embryos were used as donors for timelapse imaging, fixed at shield stages or allowed to develop at 28.5ʰC till live imaging at end of gastrulation and fixation at somite stages. Live images at shield stages were recorded using Zeiss AxioCam MRm camera; 10 hpf images were recorded using Zeiss AxioCam ICc1.

### Analysis of DNA damage in drug treated embryos

Aph+HU treated embryos were fixed at shield stages using 4% PFA. For positive control of DNA damage, dechorionated diploid embryos were irradiated with 254nm UV for 10 min at 1k-cell stage and fixed for DNA damage analysis at shield stage. DNA damage was assessed using γH2AX immunolabeling as described above.

### Differential cell labeling for live cell migration assays

For labeling small clones of cells in whole embryos, 0.5 nL of 1% Tetramethylrhodamine Dextran (Invitrogen D1817), was injected into one cell at the 16-32 cell stage in diploid, haploid or tetraploid embryos.

For co-transplanting cells to generate mosaic diploid embryos, donor diploid embryos were injected with 0.5 nL of 1% Tetramethylrhodamine Dextran and donor haploid/tetraploid embryos were injected with 0.5 nL of 1% Fluorescein Dextran (Invitrogen D1820) at the one-cell stage. At sphere stage, a few labeled cells from diploid and haploid/tetraploid donors were co-transplanted into an unlabeled diploid host embryo also at sphere stage. The initial position of the transplanted donor cells in the host embryo was recorded on a Zeiss AxioImager M2 epifluorescence microscope.

For assaying actin localization during migration, donor diploid and haploid embryos were injected at the one-cell stage with CAAX-mCherry mRNA (Addgene) to visualize membrane protrusions (Hancock, Cadwallader et al., 1991) and Alexa488-Phalloidin (0.7x in DMSO, Thermo Fisher A12379) to label actin during gastrulation. In our experiments and as has been previously reported, the concentration of Phalloidin injected did not cause developmental abnormalities (Li, Webb et al., 2008). To obtain embryos with larger cell sizes, injected diploid embryos were treated with Aph+HU as described above. At sphere stages, a few cells were transplanted from donor diploid, haploid or diploid Aph+HU embryos into unlabeled diploid host embryos.

### Timelapse imaging during gastrulation

For all timelapse experiments, embryos were screened at the shield stage for the position of labeled/transplanted cells relative to the dorsal shield. An embryo in which the dorsal shield was laterally in focus relative to the position of the labeled/transplanted cells was mounted in 0.5% low melting agarose for live timelapse imaging on Zeiss LSM880 using a 10x objective. Timelapse was recorded from shield stages to end of gastrulation and assembled on ImageJ. Cell migration was tracked using the Manual Tracking plugin in ImageJ. Tracking data was collected from 35 cells each from four diploid, haploid and tetraploid clonal timelapses and 80 cells each from four diploid-haploid and diploid-tetraploid mosaic timelapses. Statistical significance was calculated using the Mann Whitney test in GraphPad Prism 5.0.

Timelapses of actin were also recorded similarly using a 40x objective to visualize cell membrane and actin punctae during migration. From one host embryo, during shield stages to end of gastrulation, four separate clusters of cells were imaged for ∼30 mins each. Two embryos were imaged for each condition and the timelapses were assembled in ImageJ. To record the positions of actin punctae in a migrating cell, cells were viewed as circles partitioned into four quadrants and the angles subtended by the actin punctae from the center of the circle were measured in ImageJ for the duration of the timelapse. The angle values were used to make the actin position maps in Adobe Illustrator with respect to the direction of migration.

## Acknowledgements

This work was supported by an India Alliance Wellcome Trust/Department of Biotechnology Intermediate Fellowship IA/I/13/2/501042 to SN, and by the Tata Institute of Fundamental Research, Mumbai. We wish to thank Akshina Mehta for help with RNA preparations for the transcriptome analysis. We thank SN lab members for inputs.

The authors declare that they have no conflict of interest.

## Author Contributions

TM performed all experiments, compiled and analysed data. SN designed the study, performed experiments, data analysis and compilation with TM. TM, ASB and SN performed cell migration quantifications and analysis. TM and RK performed the qRT-PCR experiments. SN wrote the manuscript with comments from TM.

## Animal use ethics statement

All animal husbandry and experiments were conducted in accordance with national and institutional animal use guidelines (approval number TIFR/IAEC/2015-3).

## Data availability statement

RNA sequencing data sets generated will be deposited with GenBank upon publication and will be freely available for use.

