## Supplementary material for "Dynamically Evolving Cell Sizes During Early Development Enable Normal Gastrulation Movements In Zebrafish Embryos": Supplemetary Information

**This file includes:**

Supplementary Figures 1 to 7

Supplementary Table 1

Captions for Supplementary Movies 1 to 14

**Other supplementary materials for this manuscript include the following:**

Supplementary Movies 1 to 14

**SUPPLEMENTARY FIGURES**

**Supplementary Figure 1**

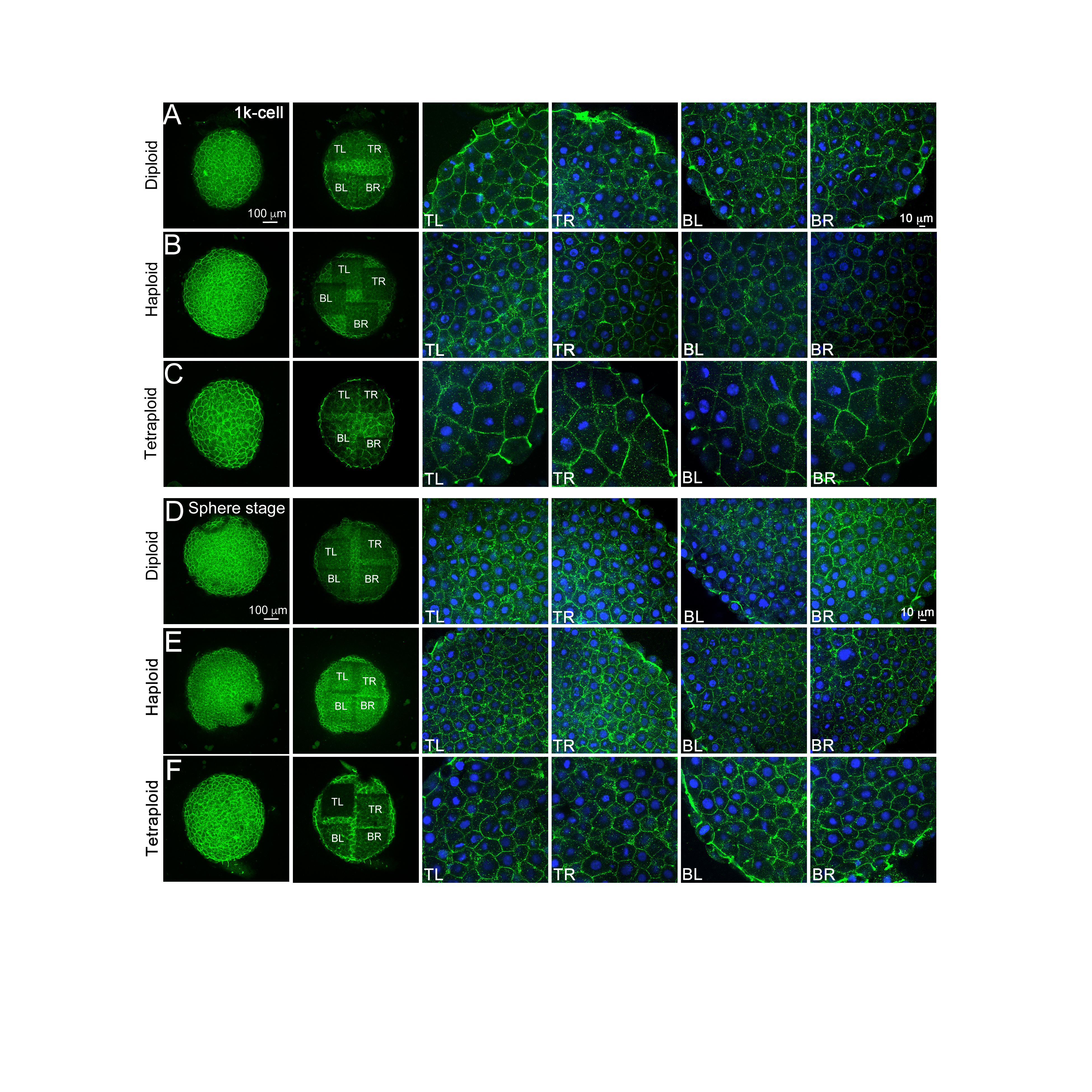

**Supplementary Figure 1: Cell sizes in diploid, haploid and tetraploid late blastoderm stages.** β-catenin immunolabels of zebrafish blastoderms (A-F). Animal view of whole blastoderms at 1k-cell (3hpf) and sphere stage (4 hpf), quadrants imaged for cell size quantifications and each of the four imaged quadrants at higher magnification (A-F). TL = top left, TR = top right, BL = bottom left and BR = bottom right.

**Supplementary Figure 2**

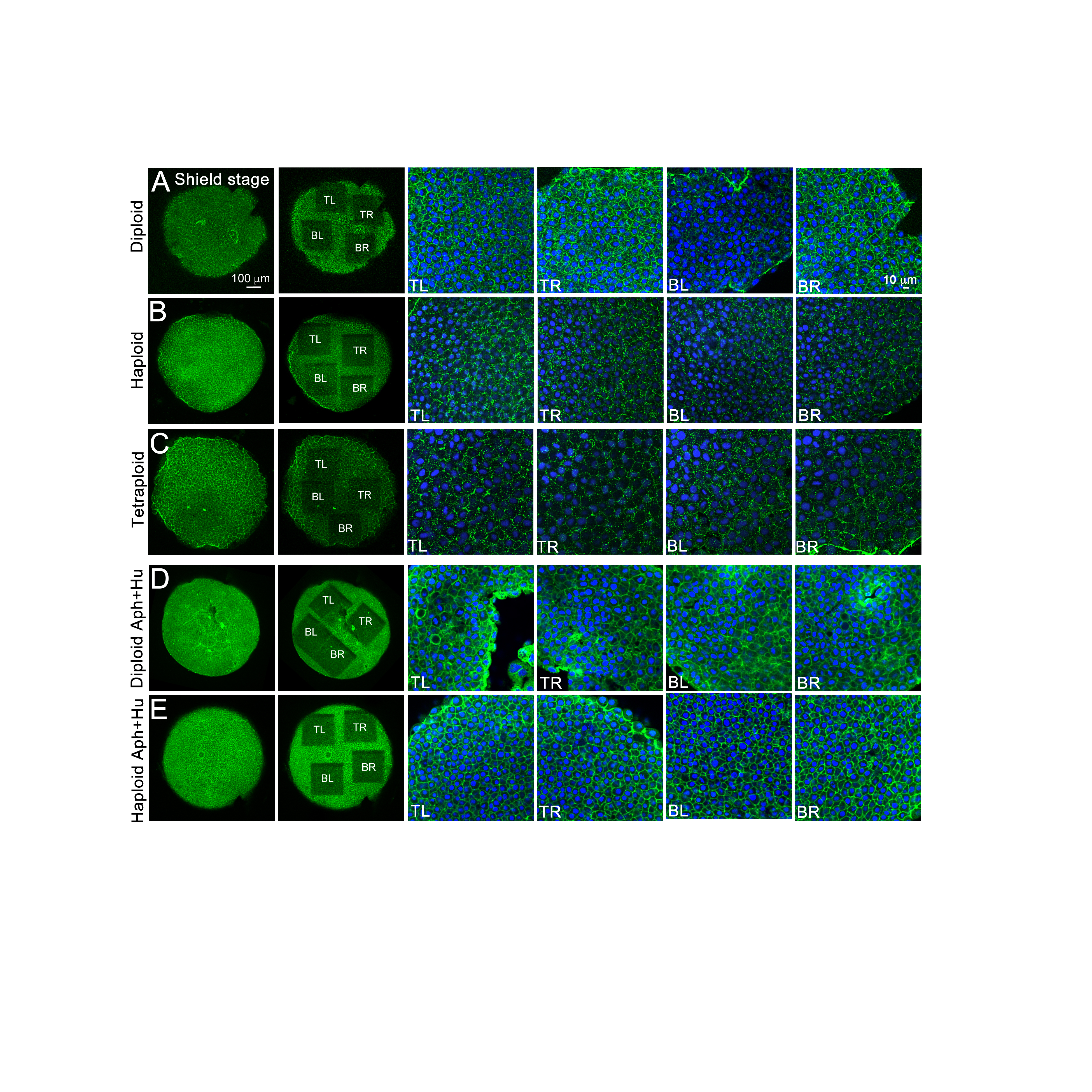

**Supplementary Figure 2: Cell sizes in embryos from different experimental conditions during early gastrula stages.** β-catenin immunolabels of zebrafish blastoderms (A-E). Animal view of whole blastoderms at shield stage (6 hpf), quadrants imaged for cell size quantifications and each of the four imaged quadrants at higher magnification (A-E). TL = top left, TR = top right, BL = bottom left and BR = bottom right.

**Supplementary Figure 3**

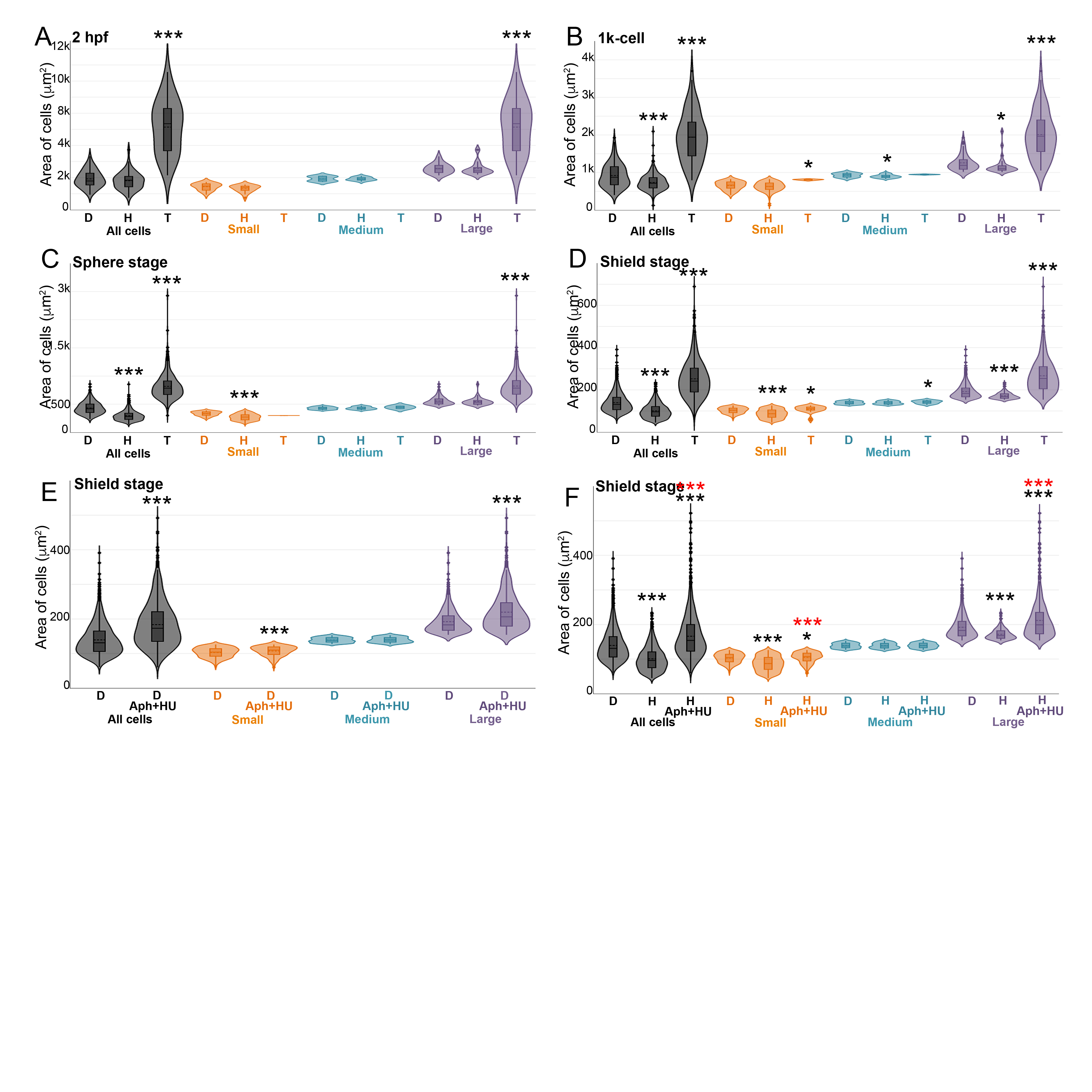

**Supplementary Figure 3: Distributions of cell areas across developmental stages.**

Cell area distributions across development from haploids and tetraploids in comparison to diploids, categorized into small, medium and large cell size bins based on diploid sizes, dotted horizontal line within box plots indicate mean area (A-D). Cell area distributions in diploid Aph+HU embryos at shield stages, binned into size categories according to diploid cell size values (E). Cell area distributions in diploid, haploid and haploid Aph+HU embryos at shield stages, binned into size categories according to diploid area values (F). Asterisks denote statistical significance based on Mann Whitney (p ≤ 0.05). In F, black and red asterisks indicate statistical significance in comparison to diploids and haploids, respectively.

**Supplementary Figure 4**

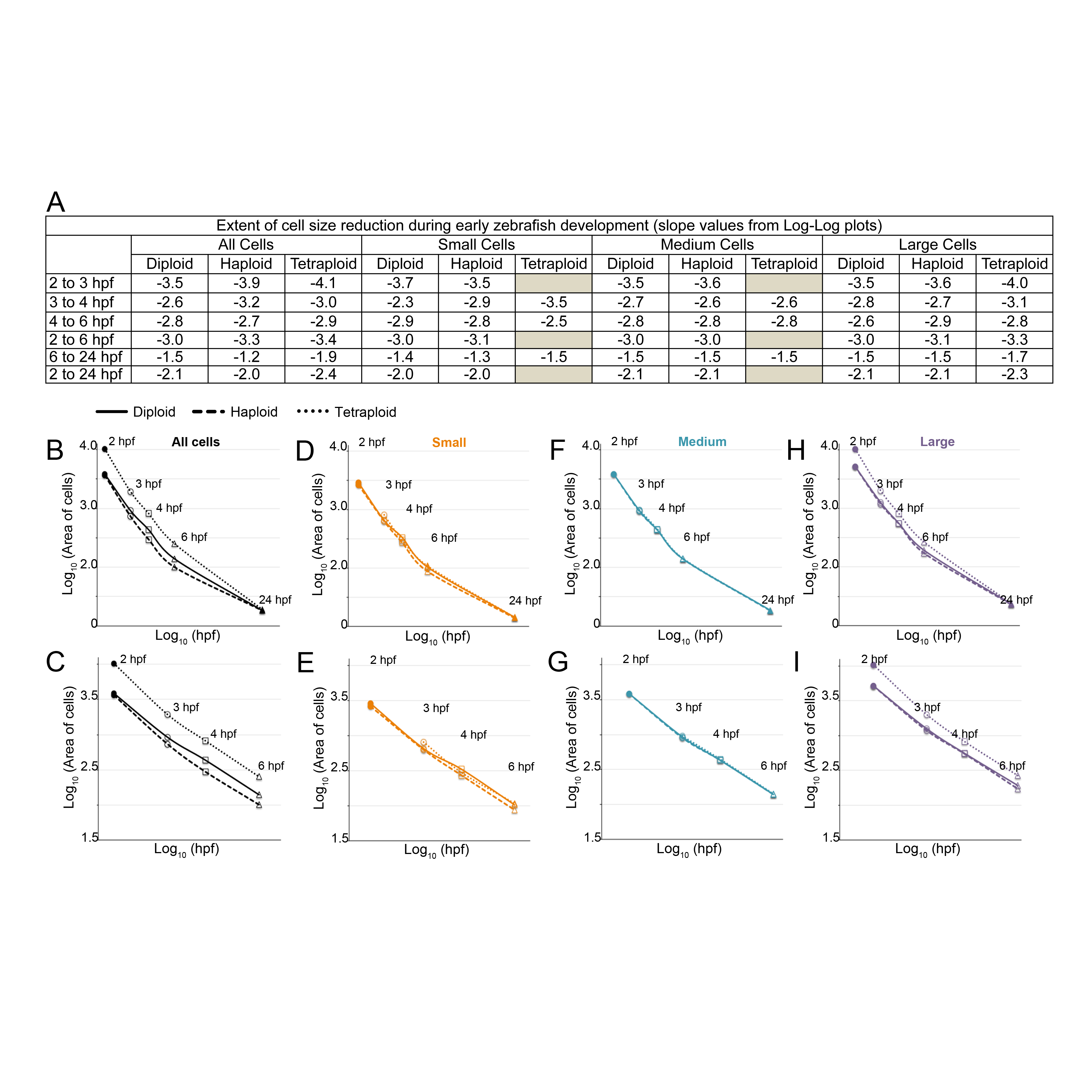

**Supplementary Figure 4: Quantitative analysis of cell size reduction during early zebrafish development and in altered ploidy embryos.** Extent of cell size reduction calculated as slope from log-log plots of absolute cell area and developmental time in diploid, haploid and tetraploid embryos from Fig.1P, Q and from panels B to I. Extent of cell size reduction in diploids (D), haploids (H) and tetraploids (T) for all cells, small, medium and large cells during development plotted as log-log plot (B-I).

**Supplementary Figure 5**

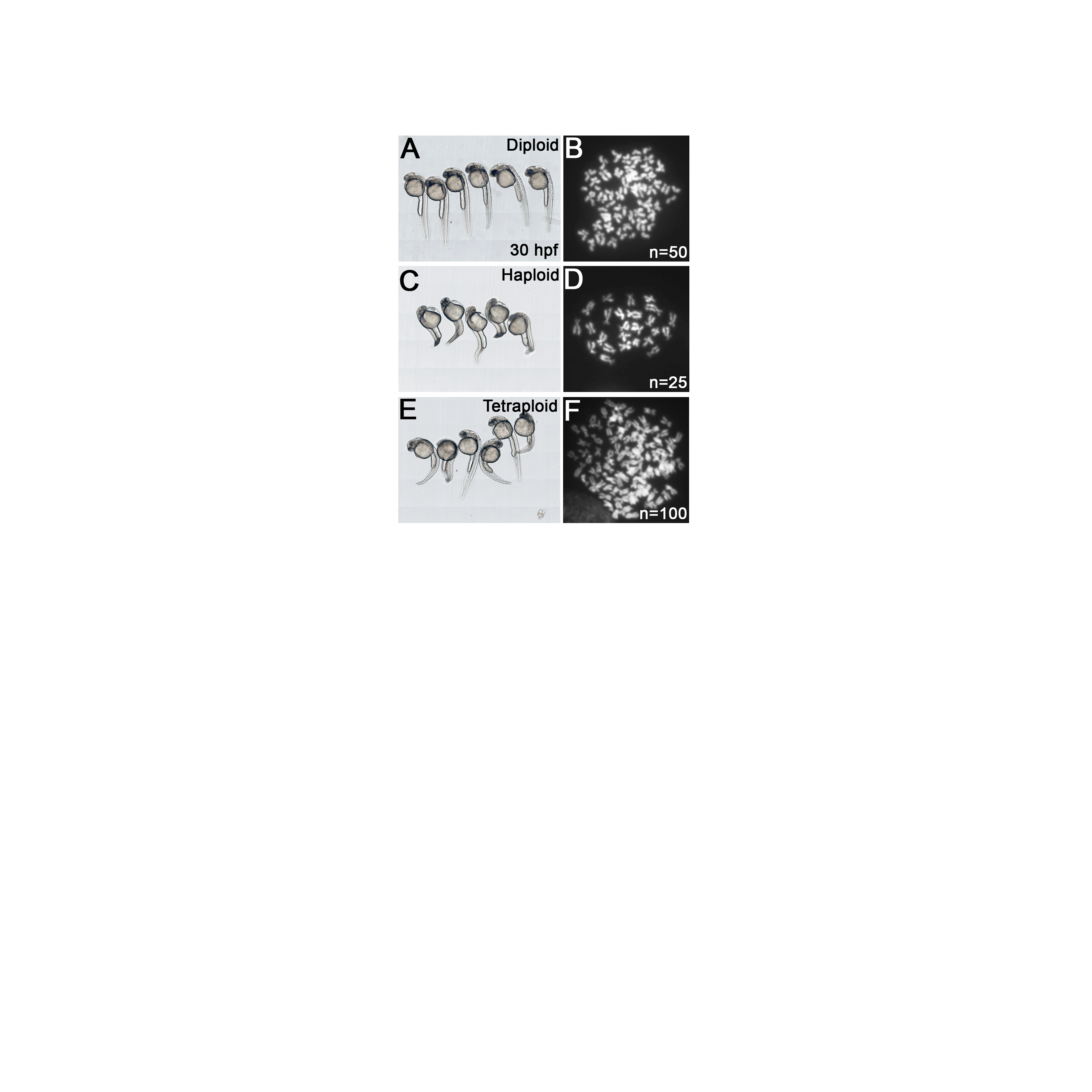

**Supplementary Figure 5: Late developmental abnormalities and metaphase chromosomes spreads.**

Images of live diploid (A), haploid (C) and tetraploid (E) embryos at ~30 hpf, which shows that haploid and tetraploid embryos are developmentally abnormal and not just delayed. Metaphase chromosome spreads show half the chromosome number in haploids (D) and double the chromosome number in tetraploids (F) in comparison to diploid (B).

**Supplementary Figure 6**

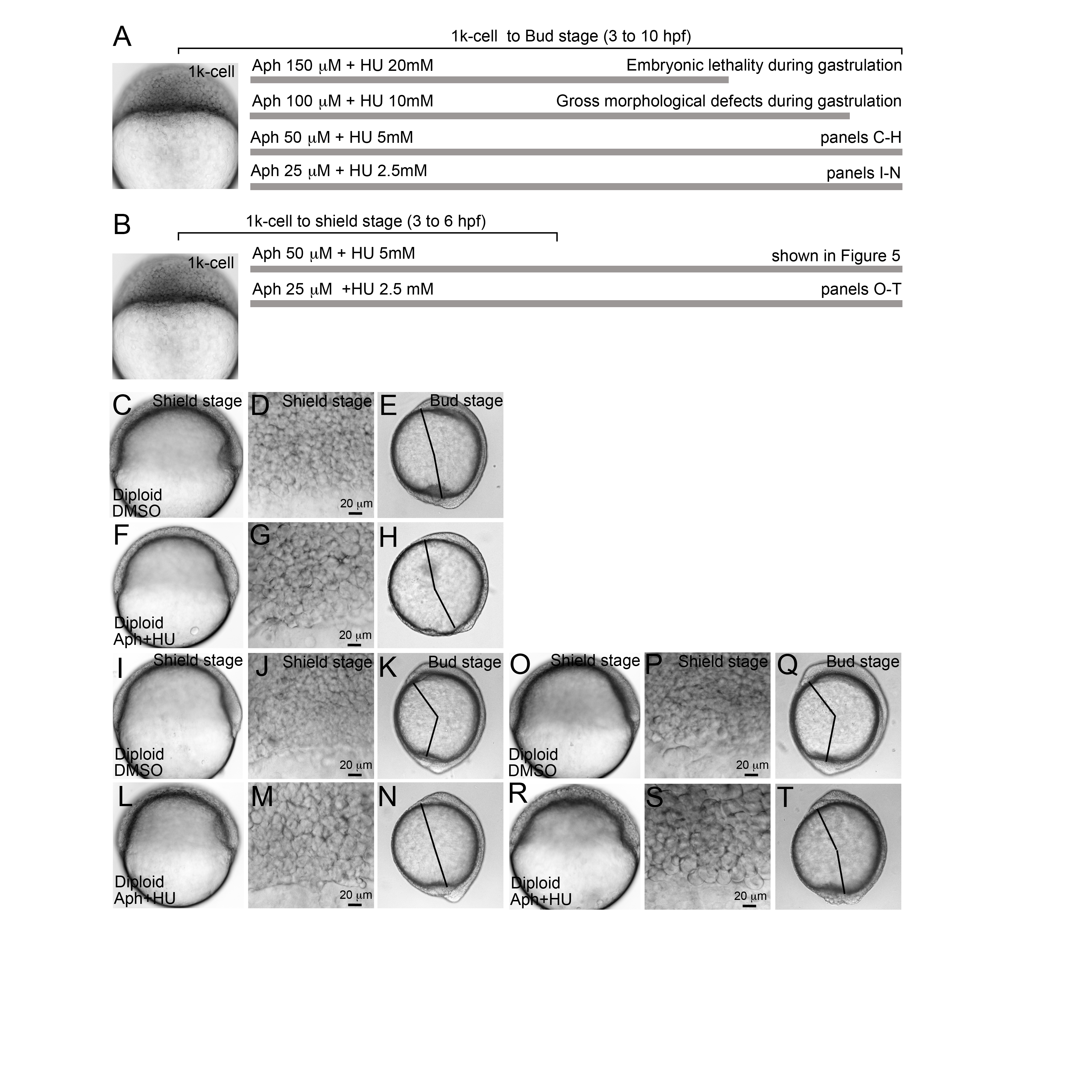

**Supplementary Figure 6: Standardization of Aphidicolin and Hydroxyurea exposure regime in diploids to alter early cell sizes before gastrulation.**

Schematic of Aphidicolin and Hydroxyurea exposures as shown in A, embryos were incubated in drug or DMSO containing medium starting at 1k-cell (3 hpf, A) to Bud stage (10 hpf) and imaged at shield stage (6 hpf) and Bud stage (C-N). By shield stages, drug treatments resulted in an increase in cell size in diploids (F, G, L, M) in comparison to DMSO treated diploid embryos (C, D, I, J). By end of gastrulation at Bud stage, drug treated diploid embryos had shorter body axis (H, N) in comparison to DMSO (E, K). Schematic of Aphidicolin and Hydroxyurea exposure as shown in B, embryos were incubated in drug or DMSO containing medium starting at 1k-cell (3 hpf, B) to shield stage (6 hpf) and imaged at shield stage and Bud stage (O-T). By shield stages, drug treatments resulted in an increase in cell size (R, S) in comparison to DMSO treated diploid embryos (O, P). By end of gastrulation, drug treated diploid embryos had shorter body axis (T) in comparison to DMSO (Q). The apparent size discrepancy in 6 and 10 hpf whole embryo images is due to the acquisition of the 6 hpf images using the Zeiss AxioCam MRm camera and the 10 hpf images using the Zeiss AxioCam ICc1.

**Supplementary Figure 7**

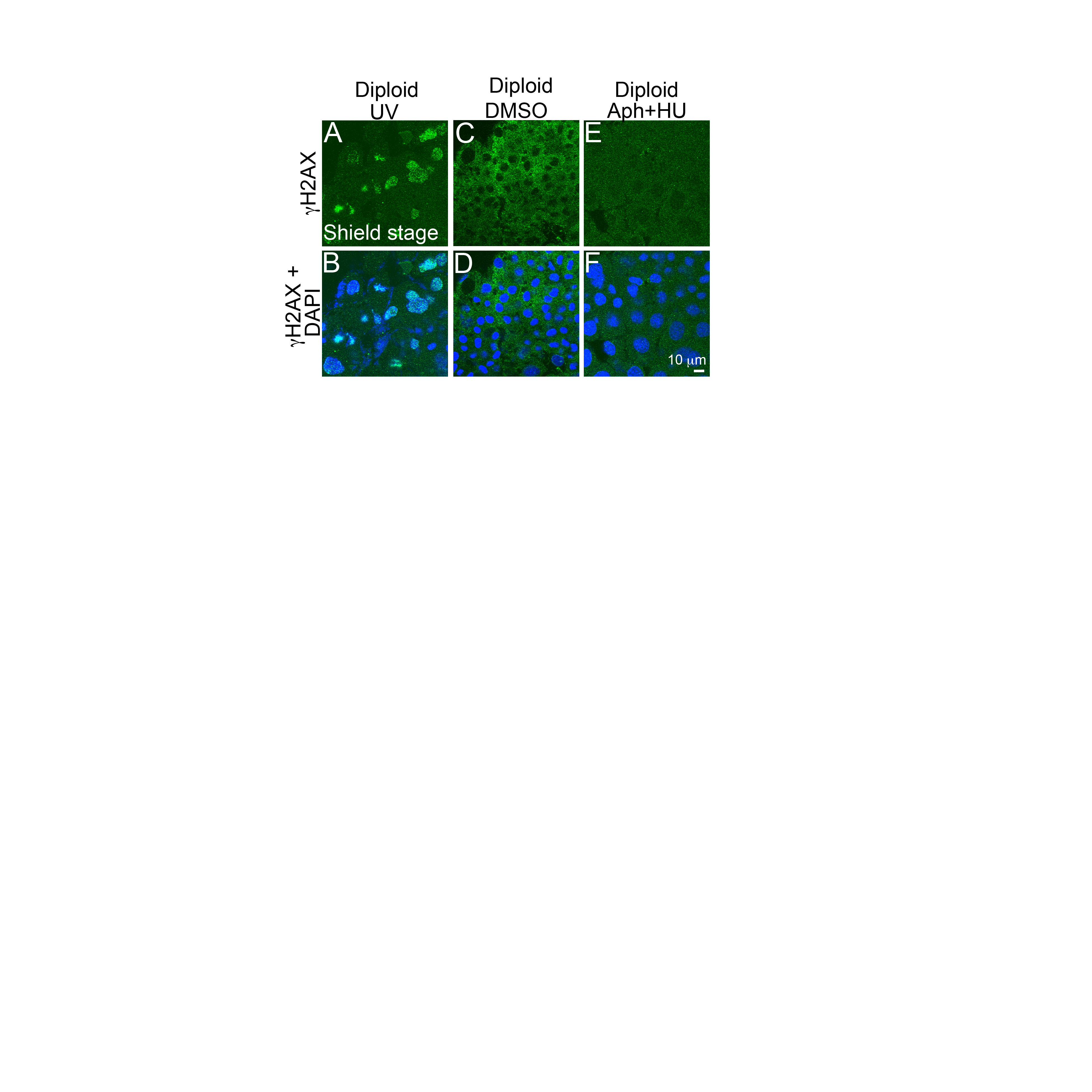

**Supplementary Figure 7: DNA damage analysis by γ-H2AX immunolabeling.** Confocal images of UV treated diploids (A, B), DMSO treated diploids (C, D) and Aphidicolin and Hydroxyurea treated diploids (E, F), stained for γH2AX (A, C, E) and with DAPI (B, D, E). Nuclei from DMSO and Aphidicolin and Hydroxyurea treated embryos do not show γH2AX staining, while those from UV-treated embryos are positive for γH2AX.

**Supplementary Table 1: Genes mis-expressed in haploids and tetraploids at 3 hpf**

Contact

**Supplementary Movie 1: Confocal timelapse of clonally labeled cells in a diploid embryo.** Injection of Rhodamine in one cell at 16-cell stage resulted in a clone of cells labeled with Rhodamine in a diploid embryo. Movie begins at ~6 hpf, with the shield oriented on the right side and is imaged through to end of gastrulation. 1 of 5 timelapse recordings is shown.

**Supplementary Movie 2: Confocal timelapse of clonally labeled cells in a haploid embryo.** Injection of Rhodamine in one cell at 16-cell stage resulted in a clone of cells labeled with Rhodamine in a haploid embryo. Movie begins at ~6 hpf, with the shield oriented on the right side and is imaged through to end of gastrulation. 1 of 5 timelapse recordings is shown.

**Supplementary Movie 3: Confocal timelapse of clonally labeled cells in a tetraploid embryo.** Injection of Rhodamine in one cell at 16-cell stage resulted in a clone of cells labeled with Rhodamine in a tetraploid embryo. Movie begins at ~6 hpf, with the shield oriented on the right side and is imaged through to end of gastrulation. 1 of 4 timelapse recordings is shown.

**Supplementary Movie 4: Confocal timelapse of co-transplanted diploid-haploid cells in a diploid host.** Timelapse imaging of co-transplanted diploid (red) and haploid (green) cells in a diploid host. Movie begins at ~6 hpf, with the shield oriented on the right-side. Haploid (green) cells have already dispersed away from the margin laterally as well as animally in comparison to diploid (red) cells. As development proceeds, red and green cells undergo epiboly and dorsal convergence towards the midline (right-side), but haploid (green) are positioned more laterally than diploid (red) cells. 1 of 7 timelapse recordings is shown.

**Supplementary Movie 5: Confocal timelapse of co-transplanted diploid-tetraploid cells in a diploid host.** Timelapse imaging of co-transplanted diploid (red) and tetraploid (green) cells in a diploid host. Movie begins at ~6 hpf, with the shield oriented on the right-side. Tetraploid (green) cells have already dispersed away from the margin laterally as well as animally in comparison to diploid (red) cells. As development proceeds, red and green cells undergo epiboly and dorsal convergence towards the midline (right-side), but tetraploid (green) are positioned more laterally than diploid (red) cells. 1 of 7 timelapse recordings is shown.

**Supplementary Movie 6: Confocal timelapse of diploid cells labeled with CAAX-mCherry and Alexa488 Phalloidin.** A heatmap of actin dynamics shown as positions of actin relative to a co-ordinate map drawn from the center of the cell of a migrating diploid cell represented as a circle. One movie spans 6 to 8 hpf of development during which a field of cells were imaged for ~30 minutes. 4 separate fields of cells were imaged from one diploid host embryo in an experiment. Two host embryos were imaged from independent experiments; movie shows one field of cells.

**Supplementary Movie 7: Confocal timelapse of diploid cells labeled with CAAX-mCherry and Alexa488 Phalloidin.** Actin and membrane mCherry labelled cells in Supplementary Movie 6.

**Supplementary Movie 8: Confocal timelapse of diploid cells labeled with CAAX-mCherry and Alexa488 Phalloidin.** Actin and membrane mCherry labelled cells merged with unlabeled diploid host cells in Supplementary Movie 6.

**Supplementary Movie 9: Confocal timelapse of haploid cells labeled with CAAX-mCherry and Alexa488 Phalloidin.** A heatmap of actin dynamics shown as positions of actin relative to a co-ordinate map drawn from the center of the cell of a migrating haploid cell represented as a circle. One movie spans 6 to 8 hpf of development during which a field of cells were imaged for ~30 minutes. 4 separate fields of cells were imaged from one diploid host embryo in an experiment. Two host embryos were imaged from independent experiments; movie shows one field of cells.

**Supplementary Movie 10: Confocal timelapse of haploid cells labeled with CAAX-mCherry and Alexa488 Phalloidin.** Actin and membrane mCherry labelled cells in Supplementary Movie 9.

**Supplementary Movie 11: Confocal timelapse of haploid cells labeled with CAAX-mCherry and Alexa488 Phalloidin.** Actin and membrane mCherry labelled cells merged with unlabeled diploid host cells in Supplementary Movie 9.

**Supplementary Movie 12: Confocal timelapse of diploid Aph+HU cells labeled with CAAX-mCherry and Alexa488 Phalloidin.** A heatmap of actin dynamics shown as positions of actin relative to a co-ordinate map drawn from the center of the cell of a migrating diploid Aph+HU cell represented as a circle. One movie spans 6 to 8 hpf of development during which a field of cells were imaged for ~30 minutes. 4 separate fields of cells were imaged from one diploid host embryo in an experiment. Two host embryos were imaged from independent experiments; movie shows one field of cells.

**Movie S13: Confocal timelapse of diploid Aph+HU cells labeled with CAAX-mCherry and Alexa488 Phalloidin.** Actin and membrane mCherry labelled cells in Supplementary Movie 12.

**Movie S14: Confocal timelapse of diploid Aph+HU cells labeled with CAAX-mCherry and Alexa488 Phalloidin.** Actin and membrane mCherry labelled cells merged with unlabeled diploid host cells in Supplementary Movie 12.
